# Increasing sensitivity and accuracy of brain-wide quantitative studies in light-sheet microscopy

**DOI:** 10.1101/230540

**Authors:** Caroline M. Müllenbroich, Ludovico Silvestri, Lapo Turrini, Tommaso Alterini, Antonino P. Di Giovanna, Irene Costantini, Ali Gheisari, Francesco Vanzi, Leonardo Sacconi, Francesco S. Pavone

## Abstract

Light-sheet microscopy (LSM) has proven a useful tool in neuroscience to image whole brains with high frame rates at cellular resolution. LSM is employed either in combination with tissue clearing to reconstruct the cyto-architecture over the entire mouse brain or with intrinsically transparent samples like zebrafish larvae for functional imaging. Inherently to LSM, however, residual opaque objects cause stripe artifacts, which obscure features of interest and, during functional imaging, modulate fluorescence variations related to neuronal activity. Here, we report how Bessel beams reduce streaking artifacts and produce high-fidelity structural data. Furthermore, using Bessel beams, we demonstrate a fivefold increase in sensitivity to calcium transients and a 20 fold increase in accuracy in the detection of activity correlations in functional imaging. Our results demonstrate the contamination of data by systematic and random errors through Gaussian illumination and furthermore quantify the increase in fidelity of such data when using Bessel beams.

## Introduction

The brain is an immensely complex entity in which structure and function are intricately correlated and best elucidated on a organ-wide scale at cellular resolution. One technique, which is particularly suited for whole-brain investigations, is light-sheet microscopy (LSM) ***(Siedentopf and Zsigmondy (1902))*** due to its intrinsic optical sectioning capabilities, fast acquisition rates and low photobleaching. In LSM, fluorescence is excited in a thin sheet of excitation light that coincides with the focal plane of a perpendicularly placed detection objective ***(Huisken et al. (2004))***. By combining advanced tissue clearing methods ***(Richardson and Lichtman (2015); Silvestri et al. (2016); Tainaka et al. (2016))*** with LSM, neuronal and vascular cyto-architecture can be reconstructed over cm-sized samples like the entire mouse brain ***(Dodt et al. (2007); Susaki et al. (2014); Pan et al. (2016))*** for quantitative structural studies. On the functional side, LSM, paired with intrinsically transparent samples like zebrafish larvae, has yielded fast volumetric calcium activation maps over the entire encephalon ***(Vladimirov et al. (2014)).***

Despite its intrinsic advantages, the very nature of uncoupled, perpendicular optical pathways for fluorescence excitation and detection entails a different set of drawbacks unique to LSM. Previously published studies (***Chen et al. (2016))*** have shown that refractive heterogeneities, always present to some extent even in extremely well clarified or intrinsically transparent samples, lead to a loss of spatial resolution and a concomitant degradation in sensitivity and contrast. Particular to LSM are dark shadows that appear whenever the fluorescence-exciting light sheet is interrupted by scattering or absorbing obstacles. At best, these dark shadows severely affect image homogeneity, at worst, they completely obscure any feature of interest in their path. Considering the increasingly large dataset sizes routinely produced in high-throughput light-sheet microscopy, high demands are placed on the automated tools to count, trace or segment the fluorescent features of interest. Consequently, background uniformity and indeed high-fidelity imaging are paramount to facilitate the extraction of meaningful insights from terabytes of data.

Furthermore, if the obstacles that obstruct the light sheet are not static, their shadows dynamically modulate the fluorescence intensity, causing an artifact here termed “flickering”. For example, haemodynamic absorption ***(Ma et al. (2016))*** of the light sheet can be particularly problematic for ***in-vivo*** Ca^2+^-imaging where the variation of fluorescence signal over time quantifies neuronal activity. We hypothesize that artifacts due to streaky shadows are non-negligible and cause loss or corruption of data obtained in light-sheet microscopy experiments. This premise is supported by a previous functional study in zebrafish which had to manually exclude neurons affected by severe flickering from further analysis ***(Panier et al. (2013)).*** We argue that dynamic flickering constitutes an inherent source of artifact potentially falsifying quantitative data obtained in light-sheet microscopy experiments and therefore any theory inferred from such observations might have to be questioned.

Aiming for an optical solution to streaking artifacts, here, we apply Bessel beams ***(Durnin et al. (1987),*** see box 1) to LSM to investigate biological samples for high-fidelity interrogation of their structure or function. Using a direct comparative analysis between Bessel and Gaussian illumination we provide supporting evidence and quantification of artifacts introduced by Gaussian illumination and further demonstrate that Bessel beams provide superior accuracy and sensitivity.

## Results

### Streaking artifacts obscure microscopic features of interest

In this section we present the effects of streaking artifacts in structural imaging aimed at obtaining cellular-resolution maps of the anatomy over the intact clarified mouse brain in two case studies, firstly targeting neurons and secondly, targeting the vasculature. It is apparent that in all images obtained using Gaussian illumination image homogeneity is strongly affected and dark shadows obscure microscopic anatomical features of interest. Notably those same identical features remain clearly visible when using Bessel beam illumination.

*Figure 1* summarizes shadowing artifacts in neuronal imaging demonstrated on an axial section (maximum intensity projection of 20 μm) of an intact Thy1-GFP-M mouse brain imaged with Gaussian illumination ***(Figure 1-a)*** using a custom-made LSM (supplementary ***Figure 6***). Dark horizontal shadows traverse each brain half and are further illustrated in an inset detailing the hippocampus ***(Figure 1-b)***. Note, that each half of the intact brain is illuminated from its respective side. By contrast, the same area acquired with Bessel beam illumination ***(Figure 1-c)*** shows improved image homogeneity and the absence of strong shadowing. In order to quantify the extent of the area affected by streaking artifacts, we calculated the line profiles obtained over the entire height of the image for Gauss and Bessel illumination respectively ***(Figure 1-d)*** and further binarized their absolute difference ***(Figure 1-e***) with respect to a user-selected threshold to obtain a pattern similar to a bar code. By superimposing this bar pattern to the original image ***(Figure 1-f)*** the percentage of 2D area affected by streaking artifacts was estimated. Averaged over a slab encompassing the entire brain over a depth of 400 μm (stitched from image stacks with step size 2 μm, threshold 5 %) we calculated that 37.5 % ± 0.2 % (standard error of mean, SEM) of the images were affected by streaking.

**Figure 1.**
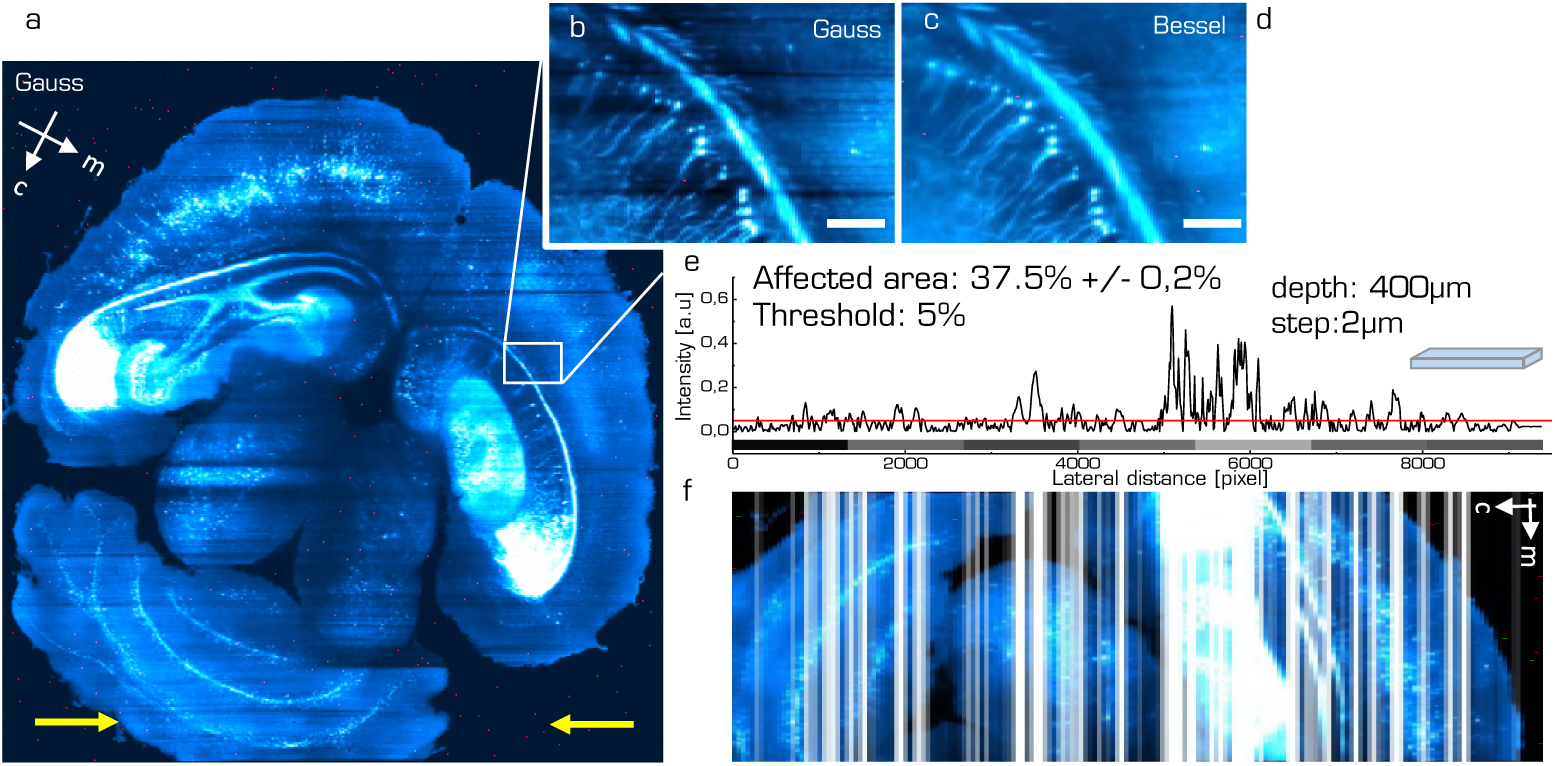
Structural neuron imaging. **(a)** Maximum intensity projection over 20 μm of a Thy1-GFP-M axial mouse brain section imaged with a Gaussian beam. Yellow arrows indicate direction of light-sheet propagation. Each half of the brain was excited by one light sheet respectively. White box marks position of details in the hippocampus affected by streaking artifacts for Gaussian **(b)** and Bessel **(c)** beam illumination respectively. Scale bar 10 μm. **d)** Line profile averaged over the entire height of the inset evidences the shadows as drops in the red curve. **e)** Calculating the absolute value of the difference in intensity line profile between Gauss and Bessel allows to estimate the area affected by streaking artifacts by applying a threshold (here 5 %). Applying this threshold to stitched images of half a brain (left half in a) **(f)** over a depth of 400 μm with a step size of 2 μm yielded that 37.5 %± 0.2 % (error is standard error of the mean, sem) of the data set was affected by streaking artifacts. **Figure 1-supplementary Figure 1**. Light-sheet microscope for structural imaging (supplementary Figure 6).

The effects of streaking artifacts on vasculature imaging are summarized in ***Figure 2*** where an axial section of mouse brain vasculature illuminated with a Gaussian beam is shown ***(Figure 2-a)***. The white box marks the position of details shown in ***Figure 2-b,c*** for Gaussian and Bessel illumination respectively. The red box marks the position corresponding to the isometric view along yz and xz illustrated in ***Figure 2-f*** for Gaussian illumination whereas the cyan box is depicted in isometric view using Bessel beam illumination in ***Figure 2-g***. Even though the whole brain data set appears to be of high quality, strong shadows in the Gaussian case completely obscure even large vessels that remain visible when illuminated with a Bessel beam (see supplementary movies 16,17). Using an automated segmentation based on simple thresholding ***(Figure 2-d,e*** and 3D projections in ***Figure 2-h,i,*** supplementary movies 18,19), the Manders coefficients were averaged throughout the stack and are reported in ***Figure 2-j***. Note, that values range from 0 to 1 and express the fraction of intensity in the Gaussian channel that is located in pixels where there is non-zero intensity in the Bessel channel and ***vice versa.*** Throughout the depth of the stack the fraction of total intensity in the Bessel channel located in pixels of non-zero intensity in the Gaussian channel was 0.62 ± 0.02 whereas the corresponding value for the Gaussian channel was 0.87 ± 0.01 (p < 0.0001, paired t-test, n=39, error is sem). Broadly speaking this signifies that while 87% of the image content present in the Gaussian channel was also present in the Bessel channel only 62% of the image content present in the Bessel channel had corresponding content in the Gaussian channel.

### Dynamic flickering falsifies functional traces

The previous section described the deleterious effect of static shadows on structural data extracted from LSM images over the entire and intact mouse brain. In the following we consider functional imaging in zebrafish where neuronal activity is quantified by a relative change in fluorescence of the calcium indicator with respect to a baseline value. Additionally to previously discussed static shadows also present in intrinsically transparent zebrafish larva ***(Figure 3-a-d***), dynamic shadowing caused by the movement of red blood cells leads to a fluctuation of this baseline, here termed flickering. The following images were acquired using a custom-built light-sheet microscope ***(Figure 7)*** which provided cellular resolution over the entire encephalon within one field of view. As early as three days post fertilization (dpf), fully formed blood vessels ***(Figure 3-e-g***, green arrow heads) can be easily distinguished by the flow of individual blood cells (red arrow heads). Haemodynamic absorption and scattering represents a multiplicative noise source ***(Ma et al. (2016))*** and therefore has the potential to modulate significantly the sensitivity to fluorescence variations related to neuronal activity in adjacent neurons (white arrow heads). A projection of the standard deviation for ≈ 100 ns of the trace ***(Figure 3-f***) clearly evidences blood vessels by superimposing the trajectories of blood cells and further highlights the strong variations in gray values in corresponding adjacent regions ***(Figure 3-g***).

**Figure 2.**
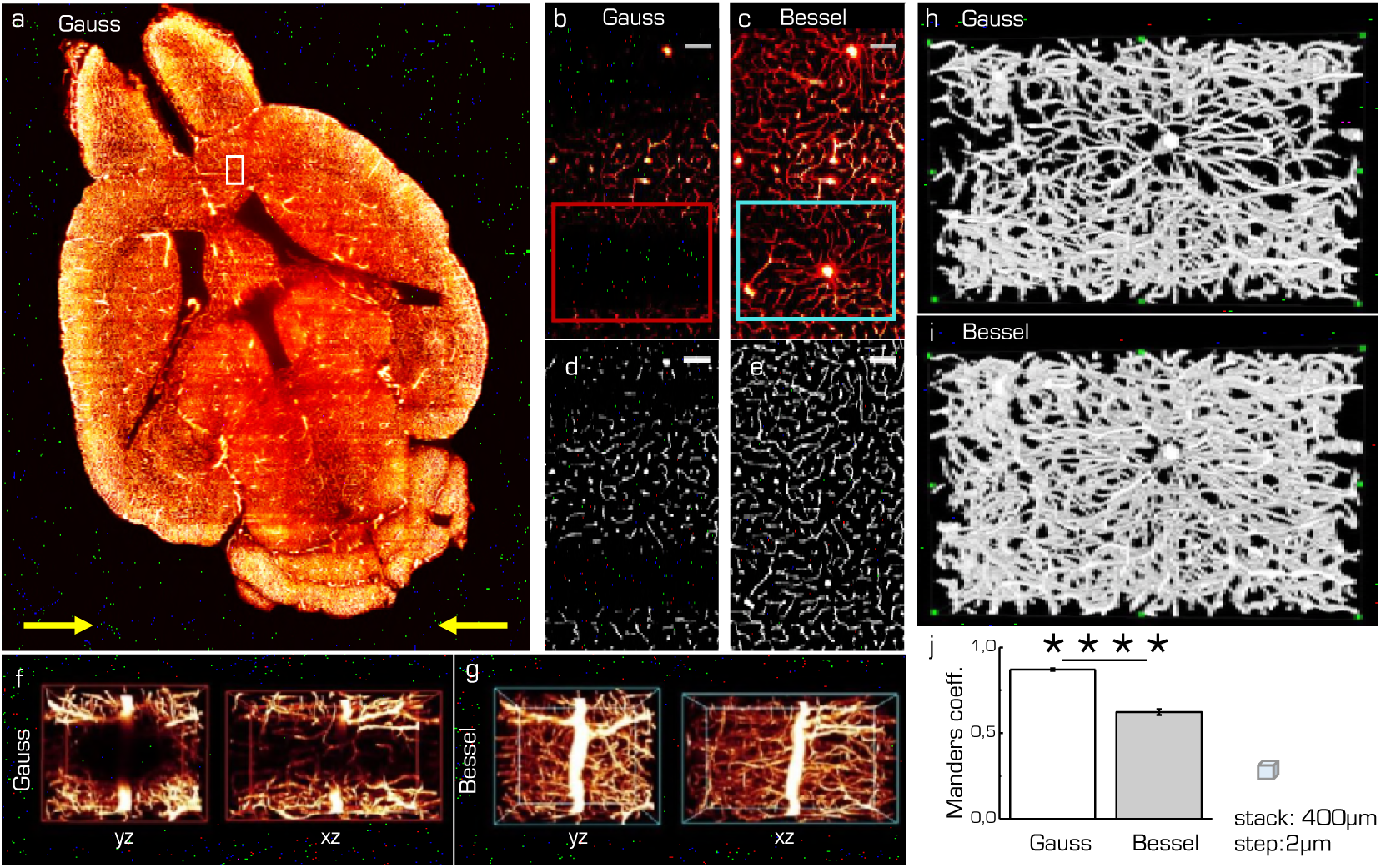
Vascular structural imaging. **(a)** Brain vasculature labeling of adult Thy1-GFP-M mice with tetramethylrhodamine-albumin imaged with Gaussian illumination. Yellow arrows indicate light-sheet propagation. White box corresponds to images (b-e). Strong shadows obscure even large vessels when illuminated with Gaussian **(b)** but not with Bessel beam illumination **(c)**. Scale bar: 10 μm. Automated segmentation based on simple thresholding for Gaussian **(d)** and Bessel beam illumination **(e). f,g)** Isometric views of red (cyan) box in b(c) shown along yz and xz for Gaussian (Bessel) illumination. **h,i**) 3D projections of the segmented data corresponding to the red and cyan ROIs indicated in (b,c). **j**) Manders coefficients averaged over a 400 μm stack comprising images in (d,e) for Gaussian and Bessel illumination. The fraction of total intensity in the Bessel channel located in pixels of non-zero intensity in the Gaussian channel was 0.62 ± 0.02 whereas the corresponding value for the Gaussian channel was 0.87 ± 0.01 (*p* < 0.0001, paired t-test, n=39, error is sem) **Figure2-supplementary Movie 1.** Raw data vasculature stack Gauss (b, supplementary Movie 12) **Figure2-supplementary Movie 2.** Raw data vasculature stack Bessel (c, supplementary Movie 13) **Figure2-supplementary Movie 3.** Segmented vasculature stack Gauss (d, supplementary Movie 14) **Figure2-supplementary Movie 4.** Segmented vasculature stack Bessel (e, supplementary Movie 15) **Figure2-supplementary Movie 5.** 3D projection vasculature Gauss (f, supplementary Movie 16) **Figure2-supplementary Movie 6.** 3D projection vasculature Bessel (g, supplementary Movie 17) **Figure2-supplementary Movie 7.** 3D projection segmented vasculature Gauss (h, supplementary Movie 18) **Figure2-supplementary Movie 8.** 3D projection segmented vasculature Bessel ((i, supplementary Movie 19))

**Figure 3.**
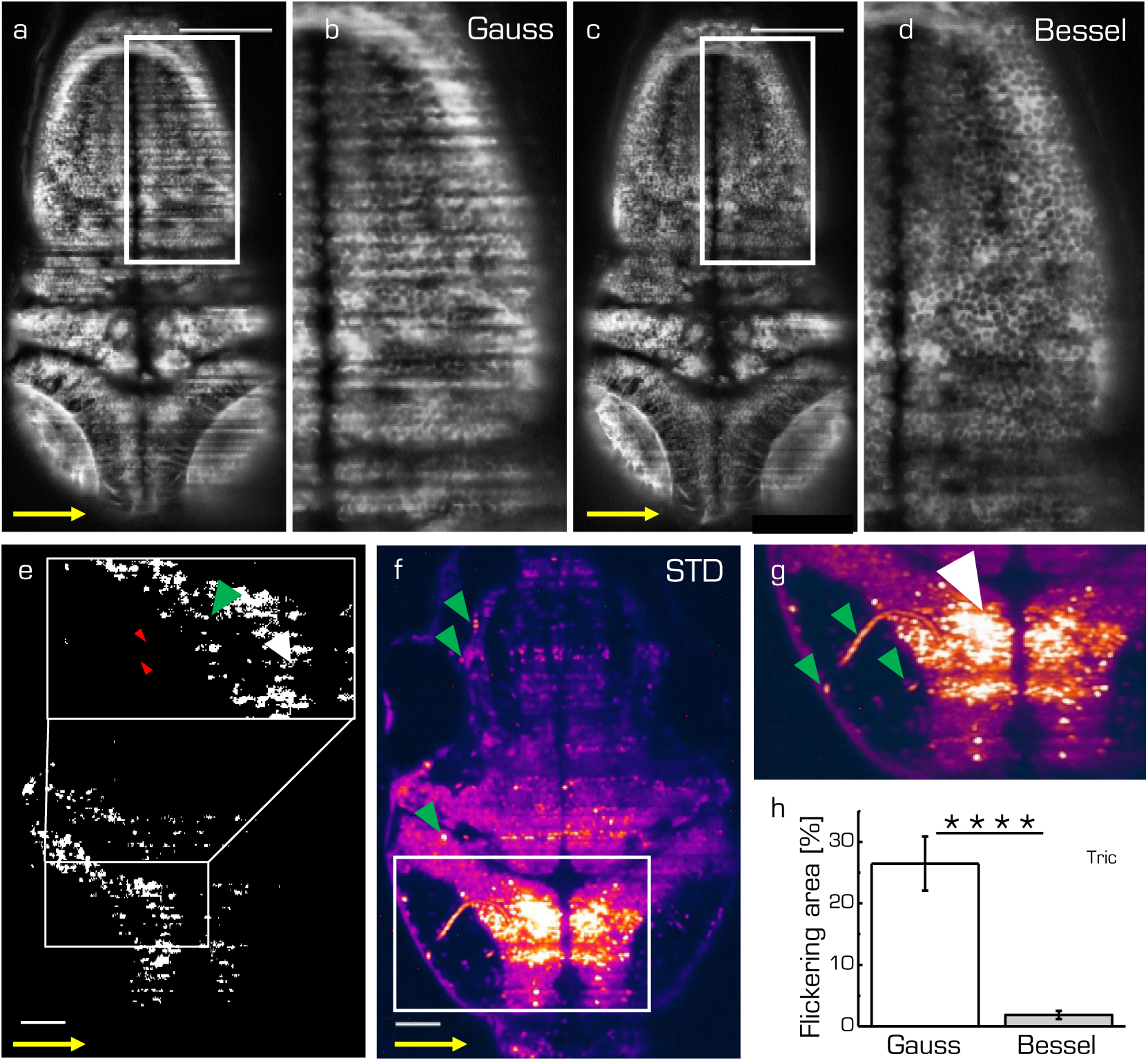
Shadow artifacts in ***in-vivo*** zebrafish imaging. **a,c)** Static shadows in the encephalon of a 3 dpf Tg(elavl3:GCaMP6s) zebrafish with cytoplasmatic expression of GCaMP imaged with Gaussian and Bessel beam illumination. Yellow arrow indicates light-sheet propagation. Scale bar: 100 μm. **b,d**) Details showing the hindbrain. e) Transverse plane of a 4dpf Zebrafish larva (Tg(elavl3:H2B-GCaMP6s)). Red blood cells (red arrow heads) passing through vasculature (green arrow head) dynamically absorb or scatter the excitation light sheet (yellow arrow) and create areas of strongly fluctuating shadow artifacts (white arrow head). Scale bar: 50 μm. **f**) Projection of the standard deviation over ≈ 100 ms of a time lapse recording on the plane shown in panel e. **g)** Inset of area indicated in f, each of the three segments of vasculature (green arrow heads) causes a corresponding area of high standard deviation of the fluorescence intensity (white arrow head). **h**) Quantification of 2D area strongly affected by flickering for Gaussian (23.8 % ± 6.5 %) and and Bessel beam illumination (0.8 % ± 0.5 %) (*p* < 0.0001, paired t-test of n=18 planes in 10 larvae aged 4-5 dpf, error is sem). Tricaine (160mgl^-1^) was added to the fish water. **Figure3-supplementary Figure 1**. Light-sheet microscope for functional imaging (supplementary Figure 7) **Figure3-supplementary Movie 1.** Raw data time lapse imaged with a Gaussian beam (supplementary movie 20) **Figure3-supplementary Movie 2.** Raw data time lapse imaged with a Bessel beam (supplementary movie 21) **Figure3-supplementary Movie 3.** Standard deviation time lapse imaged with a Gaussian beam (supplementary movie **22**) **Figure3-supplementary Movie 4.** Standard deviation time lapse imaged with a Bessel beam (supplementary movie 23) **Figure3-supplementary Figure 2.** Estimation of the flickering area (supplementary Figure 8) **Figure3-supplementary Figure 3.** Extending bounding boxes (supplementary Figure 9)

In this section we evaluate to which extent flickering potentially masks or even falsifies neuronal activity when using Gaussian illumination. Furthermore we directly compare each quantification with its Bessel beam counterpart. Based on an approach employing standard deviation projections, we devised an automated methodology detailed in the methods ***subsection*** and supplementary ***Figure 8*** and ***Figure 9.*** Tricaine, a general anesthetic that blocks voltage sensitive Na+ channels preferentially in neurons, was added to the fish water. The 2D area fraction affected by strong flickering was quantified at 23.8 % ± 6.5 % for Gaussian and 0.8 % ± 0.5 % for Bessel beam illumination respectively ***(Figure 3-h***, ***p <*** 0.0001, paired t-test, n=18 planes in N=10 larvae aged 4-5 dpf).

#### Haemodynamic baseline contamination

Due to the irregular flow of blood cells, the traversing excitation light sheet is either absorbed or scattered out of its trajectory such that the baseline fluorescence in adjacent neurons is no longer steadily generated. In the following, dF/F traces of pairwise identical neurons, imaged with either a Gaussian or Bessel beam, located in areas affected by strong flickering and located as described in the previous section, were compared. Larvae were treated with tricaine (160 mg I^-1^), a commonly used general anesthetic to globally lower neuronal activity ***(Attili and Hughes (2014); Turrini et al. (2017)),*** to ensure that any changes in baseline fluorescence were only due to dynamic flickering and not neuronal activity. A transversal section of the encephalon of a 4 dpf Tg(elavl3:H2B-GCaMP6s) zebrafish larva is shown in ***Figure 4-a,b***. Three representative neurons are marked with red (cyan) circles for the Gaussian (Bessel) case and their dF/F traces are shown in ***Figure 4-c***. Different positions in the encephalon display different levels of baseline noise; however, in general, the noise obtained in Bessel traces was consistently lower than that in traces obtained with Gaussian illumination. Quantified by their standard deviation, we analyzed traces measured with Gaussian and Bessel beam illumination respectively to obtain the level of baseline noise in the absence of neuronal activity ***(Figure 4-d***). The baseline noise was 19.65 % ± 0.18 % of dF/F for Gaussian and 3 .90 % ± 0.07 % for Bessel beam illumination (*p* < 0.0001, paired t-test, n=625 cells, 15 time lapses at various depths in N=7 larvae of 4-5 dpf, error is sem) constituting a five fold increase in sensitivity.

#### Statistical analysis of reduced sensitivity and accuracy

The previously reported increase in baseline noise directly reduces the sensitivity to calcium transients. In this section, using statistical arguments, we quantify the loss in peak detection and activity correlation due to time-varying haemodynamic flickering.

Exemplary traces in ***Figure 4-e*** illustrate a peak counting routine (see methods ***subsection***) which counted peaks above Gaussian baseline noise (red triangles) and again above Bessel beam baseline noise (cyan triangles) on identical traces obtained with Bessel beam illumination. Peaks of sub-threshold prominence (black arrow head) were discarded. The total number of peaks per minute detected above their respective baseline noise levels are reported in Figure 4-f(*p* < 0.0001, paired t-test, n=586 cells, in N=1 larva, error is sem). The cells were located at 12 different depths throughout the encephalon of a 4dpf larva. Whereas on average 6.6 peaks per minute could be detected above Bessel beam noise levels, only 0.5 peaks per minute were high enough to surpass the Gaussian baseline noise. This means that, statistically, if a given cell were to be located in an area affected by strong flickering, then less than 1 out of 12 peaks would be detected due to the higher associated baseline noise when using Gaussian compared to Bessel beam illumination.

**Figure 4.**
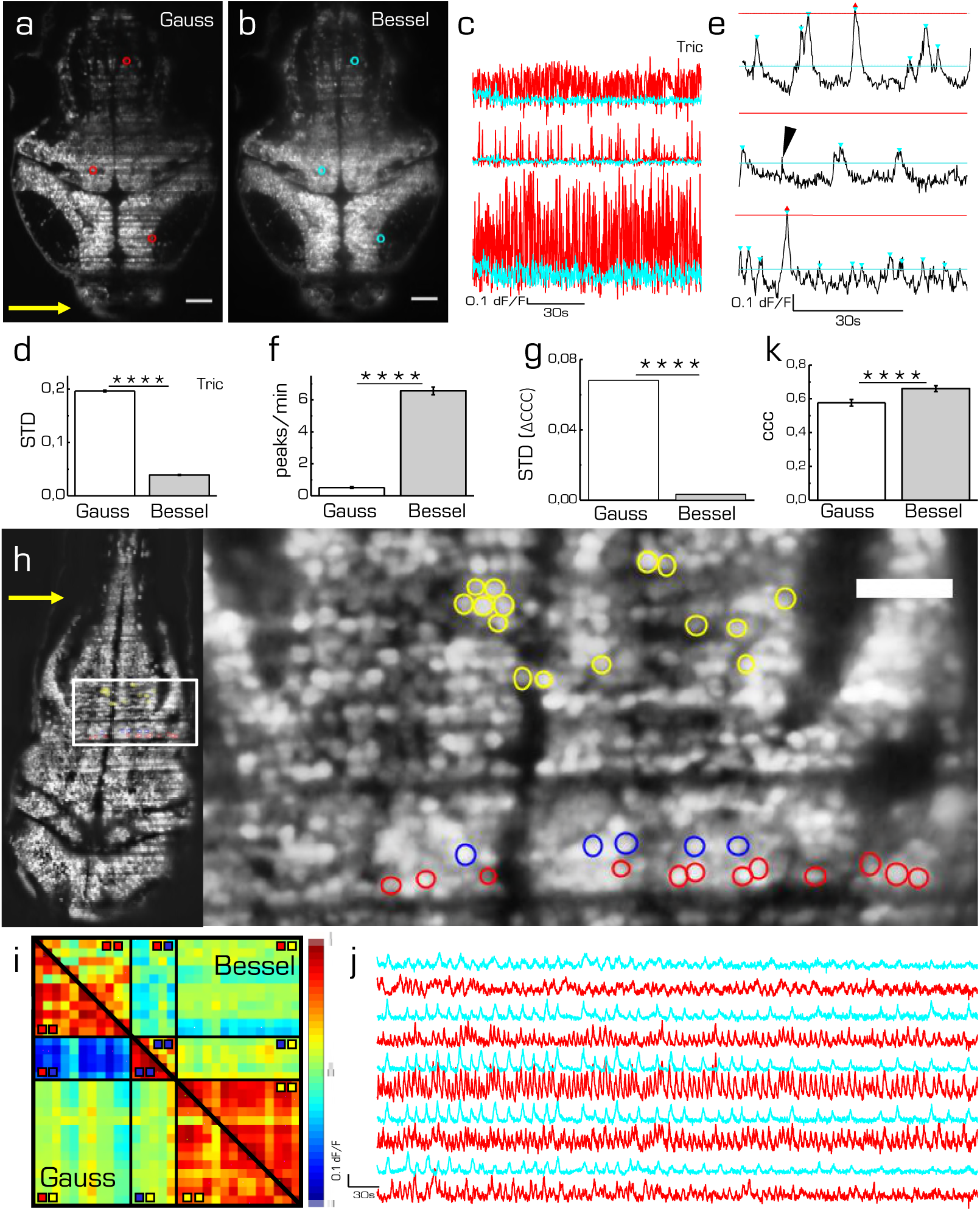
Ca^2+^-imaging in zebrafish. Transverse plane of a 4dpf elavl3:H2B-GCaMP6s larva imaged with Gaussian **(a)** and Bessel beam illumination **(b)**. Scale bar is 50 μm. Yellow arrow indicates direction of light sheet. Indicated are three exemplary cells with red (cyan) circles which have been illuminated with a Gaussian (Bessel) beam and whose traces can be seen in **c**). **d**) mean standard deviation (STD) of traces without neuronal activity (*p* < 0.0001, paired t-test, n= 625 cells, error is sem). **e)** exemplary dF/F traces measured with Bessel beam illumination during neuronal activity. Indicated are peaks (triangles) above the noise level for Gaussian (red) and Bessel beam illumination (cyan). Peaks with prominence below threshold were discarded (black arrow head). **f**) Peaks per minute detected above noise level using appropriate thresholding to exclude Gaussian and Bessel baseline noise (*p* < 0.0001, paired t-test, n=586, in N=1 larva, error is sem). **g**) STD of the difference in cross correlation coefficients between native traces and the same traces with multiplied noise. (*p* < 0.0001, paired t-test, n= 277885 corresponding to 746 cells). See supplementary Figure 10 for details. **h**) Transverse plane of a 4dpf larva imaged with Gaussian illumination. Yellow arrow indicates direction of light sheet. Indicated are cells located on two adjacent excitation lines (red, blue) and randomly in the hindbrain as part of an active network (yellow). **i**) Correlation matrix of cells measured with Gaussian (lower triangular matrix) and Bessel beam illumination (upper triangular matrix). **j**) Exemplary traces of cells (marked yellow in (i)) measured with Gaussian (red) and Bessel beam illumination (cyan). **k**) Averaged coefficient of cross correlations (ccc) of yellow yellow quadrant (*p* < 0.0001, paired t-test, n= 91, error is sem). **Figure 4-supplementary Figure 1.** Correlation with synthetic noise (supplementary Figure 10) **Figure 4-supplementary Figure 2.** Paired cross correlation coefficients (supplementary Figure 11) **Figure 4-supplementary Movie 1.** Time lapse with Gaussian illumination (supplementary Movie 24)7 of 29 **Figure 4-supplementary Movie 2.** Digital zoom onto ROIs in time lapse with Gaussian illumination (supplementary Movie 25)

To further illustrate the influence of increased baseline noise on dF/F traces, we generated random white noise of an amplitude corresponding to the Gaussian and Bessel baseline noise respectively, and multiplied it pairwise to traces obtained with Bessel beam illumination, see supplementary ***Figure 10.*** The native data set without synthetic noise was considered as ground truth and allowed an absolute comparison between Gaussian and Bessel beam illumination. The absolute cross correlation coefficients reported in supplementary ***Figure 10-e*** reveal that while the mean of the Gaussian and native data set differ significantly (0.0978 vs 0.1309), the difference between the mean of the Bessel and the native data set is not statistically significant (0.1308 vs 0.1309) (*p* < 0.0001, t-test, n= 277885 corresponding to 746 cells in N=1 larva, error is too small to be displayed). Next, the difference in cross correlation coefficients between the native traces and the traces with synthetic noises was calculated (supplementary ***Figure*** 10-f,i). The histograms of these distributions give some important insights; whereas the mean value represents the bias of each modality, the standard deviation gives a measure of its accuracy. For Gaussian illumination, we obtained a mean value of 0.0127 (supplementary ***Figure*** 10-g) whereas this value is 5.29E-5 for Bessel beam illumination (supplementary ***Figure*** 10-j). This result matters, because it indicates the introduction of a systematic error by Gaussian illumination which is approximately one tenth of the average absolute correlation coefficient. Furthermore, the standard deviations of the histograms give a direct measure of the random error introduced by each method and therefore a measure of accuracy. The standard deviation was 0.0682 for Gaussian and 0.0033 for Bessel beam illumination ***(Figure 4-g)*** which constitutes an 20-fold increase in accuracy compared to Gaussian illumination.

#### Real neuronal connectivity is masked by spurious correlation

The previous section evaluated the reduction of sensitivity from a statistical point of view by distinguishing peaks with respect to numerical thresholds and by including baseline noise of typical amplitude both obtained by averaging hundreds of cells affected by flickering. We showed that the contamination with both systematic and random errors is significantly higher when using Gaussian beam compared to Bessel beams. In the final section, we present measurements of spontaneous neuronal activity in individual cells in the presence of flickering.

As indicated in ***Figure 4-h***, several cells have been manually selected in the hindbrain of a 4dpf larva to generate a scenario in which spurious correlations due to flickering significantly alter the correlation amongst cells in a circuit. The red and blue cells where chosen in an area determined by the methodology described in ***Figure 3-h*** and corresponded to lines in which nearby flowing blood cells cast alternating shadows whereas the yellow cells have been randomly chosen among a circuit of cells that showed strong spontaneous activity (see supplementary movies 24 and 25). It is apparent from the correlation matrix ***(Figure 4-i***) that, when using Gaussian illumination, a strong correlation between cells on one line (sector marked 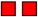 in the correlation matrix) can be observed which strongly anti-correlates with cells from the line immediately next to it 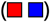, an obvious haemodynamic artifact. Notably when using Bessel beam illumination this strong spurious correlation is not observed (average cross correlation -0.571 ± 0.025 for Gauss versus -0.013 ± 0.023 for Bessel beam illumination, n=55, error is sem, see table in supplementary ***Figure 11***). Exemplary traces of spontaneous activity outside an area strongly affected by flickering (cells marked yellow) are shown in ***Figure 4-j***. It is worthwhile pointing out, that when looking at correlations entirely outside flickering areas 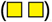, average correlation obtained with Gaussian beams was 0.576 ± 0.020 and 0.660 ± 0.017 (an increase of 15 %, n=91) for Bessel beams. This result is important because it demonstrates that even when steering clear of problematic areas with evident strong flickering, correlations of neuronal activity can be significantly affected by residual flickering when using Gaussian illumination.

## Discussion

### Structural imaging

A particular interest in the neuroscience community is to map quantitative data of the whole mouse brain onto common atlases, a task for which light-sheet microscopy is particularly well suited once the mouse brain has been appropriately rendered transparent. Due to the high frame rates obtainable with LSM, whole mouse brain data sets now routinely comprise several terabytes, a size which demands automated tools (***Frasconi et al. (2014)***; ***Peng et al. (2017))*** to count, trace or segment features of interest either to obtain cyto-architectonic information over a mouse brain-wide scale or in the emerging field of digital 3D histology ***(Torres et al. (2013))*** to provide automated interpretation of images used for quantitative diagnosis ***(Bucur et al. (2015)).*** Isolating fluorescent features of interest in a heterogeneous background places higher computational demands on the algorithms used, often with concurrent increase in computation time and complexity of the parameters to be tuned. A recent study showed that very simple algorithms like global thresholding or high pass filtering require uniform background intensity and fail to segment simple fluorescent forms like cell nuclei when a striated background simulating muscle fibers is added to the image ***(Chitalia et al. (2016)).*** As the complexity of the fluorescent feature increases, so do the demands on the algorithms tasked to isolate them. For instance, neuron tracing is a fundamental tool to understand neuronal morphology and function, however, the accurate segmentation of neurons is to date a challenging task due to their often complex arborization and the varying quality of microscope images ***(Acciai et al. (2016)).***

Another recent study compared automated segmentation of a simple synthetic interrupted tube with progressively added salt and pepper noise by a range of published algorithm and their failure to accurately trace this simulated neurite at noise levels of five percent ***(Liu et al. (2016)).*** In ***Figure 1,*** we show how a threshold of 5 % leads to more than a third of the image encompassing half a mouse brain to be affected by a striated background, putting at risk hours of microscope acquisition time, days of data post-processing and weeks of sample preparation. In ***Figure 2*** we have shown that, due to shadowing, image content, especially of finer vessels illuminated with a Gaussian beam, can drop to little above 40 % compared to data acquired with Bessel beam illumination, again jeopardizing entire terabyte-sized datasets. By contrast, the quality of the data sets obtained with Bessel beam illumination allowed, for example, the automated segmentation of blood vessels by simple thresholding in areas completely obscured by shadows using Gaussian illumination.

### Functional imaging

Large-scale neuronal recordings and their interpretation have been a paradigm shift towards understanding circuit function during behavior ***(Ahrens et al. (2012)).*** With substantial improvements in imaging technology and a concurrent increase in the size and complexity of neuronal data, pressure now shifts towards data analysis as a fundamental bottleneck for neuroscience ***(Freeman et al. (2014)).*** A common approach in large-scale imaging in zebrafish larvae is to apply an automatic segmentation algorithms which identifies ROIs associated with individual neurons ***(Panier et al. (2013); Kawashima et al. (2016))*** and extracts the df/f traces from the contained pixel values. The results presented in this paper demonstrate the contamination of data by artifacts created by haemodynamic flickering that threaten the automated extraction of calcium transients over the entire larva encephalon. Here, we have shown a reduction of the flickering area by a factor of ≈ 30 when using Bessel beam illumination. Whereas a previous publication ***(Panier et al. (2013))*** manually excluded severely affected neurons from further analysis, using Bessel beams would allow to include substantially more cells in a move from large-scale imaging towards true brain-wide analysis.

The deleterious effect of flickering artifacts caused by passing blood cells has been previously reported ***(Panier et al. (2013))*** but not quantified. Here, we present a statistical analysis to estimate the baseline noise associated haemodynamic modulation in most adversely affected neurons and report a STD of ≈ 20 % df/f for Gaussian illumination compared to ≈ 4 % for Bessel beams. This represents a 5-fold increase in sensitivity to accurately reveal Ca^2+^ transients when using Bessel beam illumination that otherwise would have been buried in noise. It is worth pointing out that these values represent a worst-case scenario; in the best case, at least in 2D studies, the plane of interest contains no or very little vessels and therefore no flickering and consequently the baseline noise in Gaussian and Bessel beam are identical. For true 3D brain-wide acquisitions however, it will be impossible to fully avoid haemodynamic contamination.

To illustrate the effect of increased baseline noise, we have presented a statistical analysis applying peak counting and cross correlation to identical traces. By counting peaks due to spontaneous neuronal activity above the Gaussian baseline noise and the Bessel baseline noise we estimated that less than 1 out of 12 peaks would be detected. This is a conservative estimate since usually higher thresholds of signal to noise ratio (SNR), e.g. Rose criterion: SNR=5 ***(Rose (1973)),*** are common. Furthermore, using synthetic white noise, we have demonstrated that the randomizing effect of haemodynamic noise leads to a loss of correlation which cannot be hoped to be recovered by clever engineering of ever brighter calcium indicators due to its multiplicative nature.

Blood flow in the brain is regulated to match neuronal demand based on activity by restricting and dilating the diameter of vessels, a phenomenon known as neurovascular coupling ***(Attwell et al. (2010)).*** Assuming Hagen-Poiseuille law of fluidic dynamics, this results in varying volumetric flow rates and therefore speeds of red blood cells. This plus other randomizing effects like vessel orientation and location means that neurons more than just a few cell diameters away from each other experience entirely uncorrelated noise. More strikingly however, the opposite is true as well: neurons which lie in close proximity on a line parallel to the excitation light will experience a baseline noise that is very strongly correlated and capable of generating spurious correlations that are not due to activity. Playing the devil‘s advocate, we “created” a correlation matrix in which the positive correlation between cells on one line and their anti-correlation with cells on a neighboring line were so strong as to mask actual correlation due to activity in an area not severely affected by flickering. This result is important because it shows that not only do Bessel beams reduce the area affected by strong flickering in which they also generate a significantly lower baseline noise. Additionally, even in areas that are not strongly affected by flickering due to immediately close by vessels, Bessel beams reveal neuronal activity that is lost when using Gaussian illumination.

In conclusion, we have illustrated how flickering can adversely affect and even falsify data extracted from LSM images. We compared the performance of Gaussian and Bessel beam illumination in structural and functional studies, covering brain-wide morphology of neuronal and vascular networks in clarified mouse brains and Ca^2+^ imaging in larval zebrafish. We have identified sources of contamination in the form of true correlation lost in a multiplicative baseline noise and spurious correlation overpowering correlation due to actual neuronal activity when using standard illumination. We have shown how the use of Bessel beams can provide an optical solution to correct for these artifacts on the microscope system side and allow for high-fidelity imaging in light-sheet microscopes. The results presented here redefine the quality standard for quantitative measurements in LSM with a single neuron sensitivity that opens up a new class of experimental studies.

## Methods and Materials

### Perfusion protocol

Brain vasculature labeling of adult Thy1-GFP-M mice was performed using the staining protocol described in ***(Tsai et al. (2009))***, but replacing fluorescein-albumin with tetramethylrhodamine-albumin in order to avoid spectral overlap with the GFP expressed by neurons. After deep anesthesia with isoflurane inhalation, the mice were transcardially perfused with 20 ml to 30 ml of 0.01 M of phosphate buffered saline (PBS) solution (pH 7.6) and then with 60 ml of 4 % (w/v) paraformaldehyde (PFA) in PBS. This was followed by perfusion with 10 ml of a fluorescent gel perfusate, with the body of the mouse tilted by 30°, head down, to ensure that the large surface vessels remained filled with the gel. The body of the mouse was submerged in ice water, with the heart clamped, to rapidly cool and solidify the gel as the final portion of the gel perfusate was pushed through. The brain was carefully extracted to avoid damage to pial vessels after 30 min of cooling and incubated overnight in 4 % PFA in PBS at 4 °C. The day after the brain was rinsed 3 times in PBS. All experimental protocols were designed in accordance with Italian laws and were approved by the Italian Minister of Health (authorization n. 790/2016-PR).

### Gel preparation

The gel was prepared according to ***(Tsai et al. (2009))***, but replacing fluorescein-albumin with tetramethylrhodamine-albumin (no. A23016, Thermo Fisher Scientific). First, a 2 % (w/v) of porcine skin gelatin type A (no. G1890; Sigma) was prepared in boiling PBS and allowed to cool to <50 °C. The gelatin was then combined with 0.05 % (w/v) of tetramethylrhodamine-albumin (Di Giovanna ***et al.,*** manuscript in preparation). The solution was maintained at 40 °C while stirring before the perfusion.

### Whole brain clearing procedure

Mouse brains were cleared using the CLARITY technique ***(Chung et al. (2013)),*** applying the passive clearing procedure. Fixed mouse brains were incubated in hydrogel solution (4 % (wt/vol) acrylamide, 0 .05 % (wt/vol) bis-acrylamide, 0.25 % (wt/vol) VA044) in 0.01 M PBS at 4 °C for 1 week. Samples were degassed and incubated at 37 °C for 3 h to allow hydrogel polymerization. Subsequently, the brains were extracted from the polymerized gel and incubated in clearing solution (sodium borate buffer 200 nM, 4 %(wt/vol) Sodium dodecyl sulfate) (pH 8.5) at 37°C for one month while gently shaking. The samples were then washed with PBST (0.1 % TritonX in 1X PBS) twice for 24 h each at room temperature. CLARITY-processed mouse brains were optically cleared using 2,2‘-thiodiethanol (TDE), as described in ***(Costantini et al. (2015)).*** After PBST washing, the brains were serially incubated in 50 nl of 30 % and 63 % (vol/vol) 2,2‘-thiodiethanol in 0.01 M PBS (TDE/PBS), each for 1 day at 37 °C while gently shaking. After TDE clearing the brains were ready for imaging.

### *Ex-vivo* light-sheet microscope

A custom-made light-sheet microscope, specifically designed for the imaging of whole clarified mouse brains ***(Müllenbroich et al. (2015)),*** was used for all structural imaging experiments (see supplementary ***Figure* 6**). In essence, the microscope is equipped with an alternative excitation light path using two flip mirrors which directed the light towards an axicon (AX252-A, ***a*** = 2°, Thorlabs, Newton, USA) to generate a Bessel beam. The Bessel beam was filtered in a Fourier plane with a circular spatial filter to eliminate any residual Gaussian contribution. The Gaussian or Bessel beam respectively is split by a polarizing beam splitter and subsequently scanned by two galvo mirrors (6220H, Cambridge Technology, Bedford, USA) to create a double-sided illumination light sheet. The excitation objectives (Plan Fluor EPI, 10x, 0.3NA, WD 17.5 mm, Nikon, Tokyo, Japan), covered with a protective coverslip, project the light sheet into the focal plane of the perpendicularly placed detection objective (XLPLN10XSVMP, 10x,0.6NA, WD 12mm, Olympus, Tokyo, Japan) specifically designed for immersion in high-refractive index media and featured a correction collar for the refractive index of the immersion solution, ranging from 1.33 to 1.52. A tube lens forms an image onto the sensor of a fast sCMOS camera (Orca Flash4.0 v2.0, Hamamatsu, Hamamatsu city, Japan) whose line by line readout is synchronized to each step of the galvo mirrors to achieve confocal line detection. Appropriate filters are used to bandpass filter the fluorescence and reject excitation light. For vasculature imaging, excitation was λ= 561 nm and an acousto-optic tunable filter (AOTFnC-400.650-TN, AA Opto-Electronic, France) was used to regulate laser power. The samples were place in a quartz cuvette (3/Q/15/TW, Starna Scientific, Hainault, United Kingdom) containing the mounting medium (n=1.45, 63 % TDE in PBS) and placed in a custom-made chamber filled with the same mounting medium. The samples were mounted on a high-accuracy, motorized x-, y-, z-, Θ-stage (M-122.2DD and M-116.DG, Physik Instrumente, Karlsruhe, Germany) which allowed free 3D motion and rotation. The microscope was controlled via custom software written in LabVIEW 2012 (National Instruments, Austin, USA) using the Murmex library (Distrio, Amsterdam, The Netherlands).

### Image stitching and analysis

The LSM for structural imaging produces a series of 3D stacks with regions of superimpositions. To achieve a 3D image of whole specimens from raw data the Terastitcher tool ***(Bria and Iannello (2012))*** a software capable of dealing with teravoxel-sized images, was used. Graphs and data analysis were done with OriginPro 9.0 (OriginLab Corporation). Image stacks were analyzed using both Fiji (http://ji.sc/Fiji) and Amira 5.3 (Visage Imaging) software. 3D renderings of stitched images were produced from downsampled files using the Amira Voltex function. The Filament Editor of Amira was used to manually trace vessels segments. Automatic segmentation of vasculature stacks was performed with Fiji by first aligning and registering both stacks to each other using the rigid body modality in the StackReg plugin. A gamma of 1.1 was applied to both stacks over the entire image to enhance dimmer structures and an auto threshold using IsoData was applied. The Manders coefficients were calculated using the JACoP plugin.

### *In-vivo* light-sheet microscope

A custom-made light-sheet microscope ***(Figure 7-a)***, specifically designed for ***in-vivo*** imaging of zebrafish larva, was used for all functional imaging experiment. In brief, a 488 nm continuous-wave diode-pump solid state laser (Excelsior, Spectra Physics, Santa Clara, USA) with 3 mW output power was used as light source and an acousto-optical tunable filter (AOTF) was used as a shutter and power regulator. Two different illumination paths could be selected by the use of flip mirrors. The first illumination path, expands the output of the laser onto a galvanometric mirror (galvo, 6220H, Cambridge Technology, Bedford, USA). The galvo is re-imaged with a 4f telescope onto the back aperture of the excitation objective (4x, 0.13 NA, air immersion, Olympus, Tokyo, Japan). The excitation objective focused the illumination beam into the sample chamber and by applying a sawtooth waveform to the galvo the beam was rapidly scanned to form a static light sheet which coincided with the focal plane of a perpendicularly placed detection objective (20x, 1 NA, water immersion, Olympus, Tokyo, Japan). The sample was mounted on an x-, y-, z-, Θ-stage (M-122.2DD and M-116.DG, Physik Instrumente, Karlsruhe, Germany) which allowed its precise positioning and the acquisition of z-stacks. Fluorescence was detected in a wide-field scheme with a tube lens forming an image onto the chip of a sCMOS camera (OrcaFlash4.0, Hamamatsu, Hamamatsu city, Japan). Appropriate filters are used to bandpass filter the fluorescence and reject excitation light. The camera was operated in rolling shutter mode thereby creating a virtual confocal slit. In the second illumination path a telescope adjusted the beam diameter of the Gaussian beam before the axicon (AX251-A, ***α*** = 1°, Thorlabs, Newton, USA) to minimize the Gaussian contribution produced by the round tip. The self-reconstruction length of the resulting Bessel beam was then re-imaged with 3 additional lenses, each placed at 2f from each other respectively, into the common Gaussian light path. A circular spatial filter was used close to a Fourier plane near the galvo to filter out any residual Gaussian contribution to the Bessel beam. A heating system kept the sample chamber at a constant temperature of 28.5 °C. The microscope was controlled via custom software written in LabVIEW 2012 (National Instruments, Austin, USA) using the Murmex library (Distrio, Amsterdam, The Netherlands).

### Zebrafish husbandry and larva mounting

We generated two stable zebrafish transgenic lines using a Tol2 construct with elavl3 promoter that drives the expression of the genetically encoded calcium indicator GCaMP**6**s in all neurons ***(Freeman et al. (2014)).*** Tg(elavl3:GCaMP6s) line showed a cytoplasmic expression of the transgene whereas line Tg(elavl3:H2B-GCaMP6s) expressed GCaMP**6**s in fusion with histone H2B and consequently showed nuclear localization. Larvae were kept according to standard procedures ***(Westerfield (1995))***. Each 4-5dpf Tg(elavl3:H2B-GCaMP6s) larva used in the experiments was transferred into a reaction tube containing 1.5 % low gelling temperature agarose (A9414, Sigma) in fish water (150 mg Instant Ocean, 6.9 mg NaH_**2**_PO_4_, 12.5 mg Na_**2**_HPO_**4**_ per 1 l of dH_**2**_O), kept at 38 °C. The zebrafsh larva was then drawn with a syringe into a glass capillary (O.D. 1.5 mm); after gel polymerization, the agarose cylinder containing the larva was gently extruded until the head protruded from the glass ***(Figure*** 7-b). The glass capillary was then mounted onto an x-, y-, z-, Θ-stage (M-122.2DD and M-116.DG, Physik Instrumente, Germany) and immediately immersed into the sample chamber containing fish water. The fish water was kept at a constant temperature of 28.5 °C.

### Calcium imaging

For ***in-vivo*** Ca^2+^ imaging a line exposure time of 0.3 ms was used, resulting in a frame rate of 44 Hz. When using Bessel beam illumination, power was increased by a factor of three compared to Gaussian illumination. The 16bit depths images constituting each time lapse were first converted from the proprietary Hamamatsu file format to a multi-page tiff and simultaneously temporally down sampled by a factor of **8** to save disc space resulting in an effective frame rate of 5.5 Hz. When tricaine was added to the fish water and the traces therefore were used to analyze flickering, no temporal downsampling was performed. Z profiles of regions of interests were then extracted with a FIJI macro over a hand selected circular area covering the cell nucleus. The Δf/f traces were obtained with a custom-written Matlab script in which, firstly, the background was subtracted and secondly, the baseline of the traces were calculated using the ***msbackadj*** function. Thirdly, the baseline was subtracted from the trace and finally, the trace was divided by the baseline to normalize it ***(Freeman et al. (2014)).***

### Estimation of the area affected by flickering

The area affected by primary flickering for Gaussian and Bessel beam illumination was estimated in a two dimensional plane by using a sequence of custom-made macros (FIJI) and programs (LabVIEW). By primary flickering is intended strong variations in baseline fluorescence intensity directly attributable to a large, nearby blood vessel. The more subtle, yet noticeable, effect of smaller peripheral blood vessels was neglected as it was much harder to quantify. Using this automated method 2D maps of severe flickering areas were obtained which allowed us to calculate the percentage of the entire encephalon that were prone to a substantial source of additive noise. To assure that the evidenced changes in pixel brightness were not due to activity, Tricaine (160 mg l^−1^), a general anesthetic that blocks voltage sensitive Na+ channels preferentially in neurons, was added to the fish water.

The methodology ***(Figure 8***), based predominantly on a temporally down-sampled time lapse in which each frame corresponded to the standard deviation (STD) of a certain frame window of the original raw data, evidenced the characteristic shape of long horizontal stripes. A custom-written FIJI macro was used to calculate the STD time lapse of the raw data time lapse with window size 5 and step size 5. After adjusting both time lapses to the same brightness scale, an additional macro automatically compared the STD time lapses of the Gaussian and Bessel beam by running through a sequence of functions to enhance the features of the images that changed most in brightness value using the exact same parameters in both cases. With both time lapses displayed on the same brightness scale, first a gamma of 1.1, second a bandpass filter (settings from FIJI:) and third a variance filter (settings from FIJI:) were applied over the entire image. After again adjusting both time lapses to the same brightness scale, each time lapse was collapsed to a single image using using a z projection of the STD. The resulting images were thresholded to a common value and converted to 8 bit binary where dark spots indicating the parts of the encephalon that had changed the most during high frame rate (44 Hz) acquisition. The binary images where then analyzed in a custom-written LabVIEW program that automatically found the bounding boxes of each identified particle (LabVIEW function and settings:) (***Figure 9***). In the case of Gaussian illumination, the horizontal dimension of the bounding box of each particle was extended up to a mask corresponding to the free-hand selection of the encephalon of the larva. In the case of the Bessel beam illumination, the bounding box was extended to a length corresponding to the theoretically calculated reconstruction length of the Bessel beam which follows from simple trigonometric considerations:

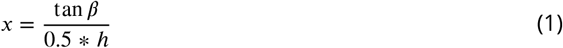

where ***β*** is the angle of the cone in the sample chamber, and ***h*** is the height of the bounding box. The LabVIEW program then automatically calculated the sum of all bounding boxes and their percentile of the free-hand selection of the entire encephalon. The characteristic horizontal stripes of the resulting areas, confirmed that the method indicated primarily shadowing artifacts and not individual neuronal activity.

### Peak counting and correlation analysis

For peak counting, the Δf/f traces were loaded into a custom-written Matlab program and smoothed using the ***sgolayfilt*** function (order 5, framelen 7). The ***findpeaks*** function was then used to automatically detect peaks above the Gaussian and Bessel noise level respectively with a minimum peak prominence of 8.5 % of Δf/f. The correlation matrices were calculated using a custom-written Matlab program using the ***corr2*** function. Noise was generated according to the following pseudo Matlab code to produce random noise that oscillates around an average of unity:

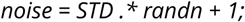

The noise vector was than point-wise multiplied to the trace vector which was averaged to zero using:

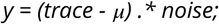

## Acknowledgments

The authors would like to thank Riccardo Ballerini and Ahmed Hajeb from the mechanical workshop at LENS for the production of custom pieces and advise on structural stability in sample mounting. The authors are grateful to Mauro Giuntini and Marco De Pas from the electronic workshop for the construction of the sample heating and electronics in the light-sheet microscopes. We further thank Misha Ahrens for providing the tol2-elavl3-H2B GCaMP**6**s and the tol2-elavl3 GCaMP**6**s plasmids (Addgene plasmid #59530 and #59531).

This project received funding from the European Union‘s H2020 research and innovation programme under grant agreements No. 720270 (Human Brain Project) and 654148 (Laserlab-Europe), and from the EU programme H2020 EXCELLENT SCIENCE - European Research Council (ERC) under grant agreement ID n.692943 (BrainBIT). The project has also been supported by the Italian Ministry for Education, University, and Research in the framework of the Flagship Project NanoMAX and of Eurobioimaging Italian Nodes (ESFRI research infrastructure), and by “Ente Cassa di Risparmio di Firenze” (private foundation).

### Box 1. Bessel beams

The paraxial Helmholtz equation governs diffraction phenomena and, most typically, its solution describes the propagation of electromagnetic waves in the form of Gaussian beams. However, alternative solutions exist in the form of diffraction-free modes, most notably Bessel beams ***(Durnin et al. (1987))***, whose central core can be extremely narrow without being subject to diffraction (***McGloin and Dholakia (2005))***. Due to finite apertures necessarily limiting the realization of such beams experimentally, only Quasi-Bessel beams can be generated for which the propagation invariance holds true over a finite propagation range ***(Durnin et al. (1988)).*** A simple way to create a Bessel beam is by superimposing plane waves whose wave vectors lie on a cone using a conical lens ***(Indebetouw (1989),*** Figure ***Box***5-a).

**Figure 5.**
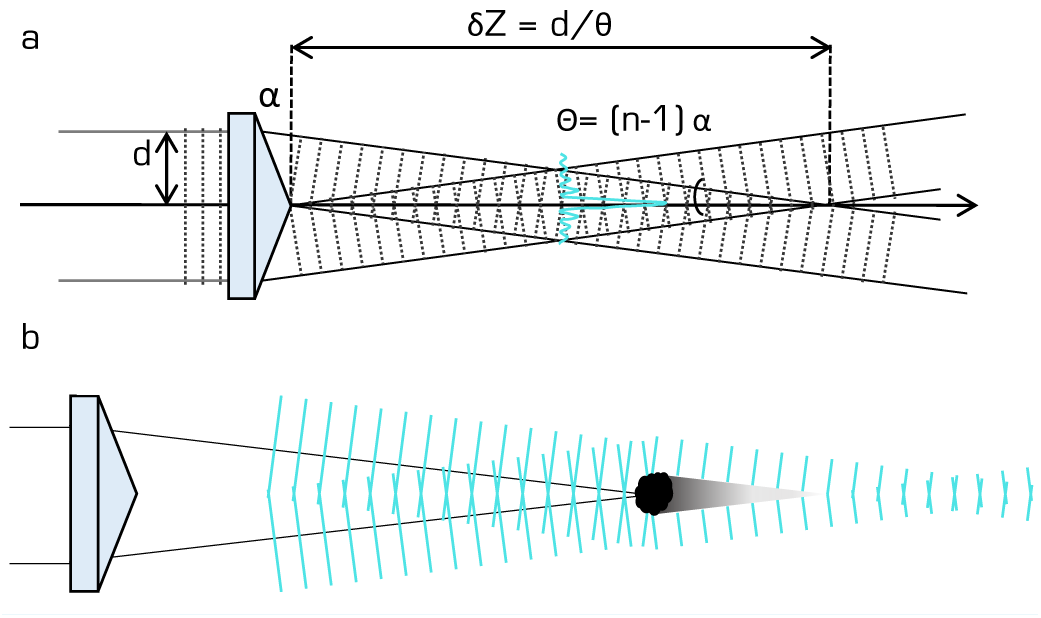
a) Generation of a Bessel beam using an axicon lens. The characteristic transverse ***J_0_*** Bessel beam profile of a central lobe with concentric rings is created within the propagation length ***δZ*** of the axicon. b) If an obstacle obstructs the central lobe of the Bessel beam, the optical power stored in the concentric rings can regenerate the initial beam^554^ profile in the reconstruction region behind the shadow zone.

The axicon creates the characteristic circularly symmetric beam profile of a tight central core with concentric rings within its depth of focus ***δZ*** given by:

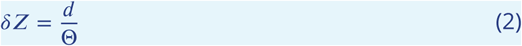

where ***d*** is the radius of the Gaussian beam impinging on the axicon and Θ is the Bessel beam cone aperture depending on the refractive index ***n*** and angle of the conical lens α:

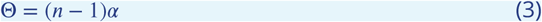

Each concentric ring carries approximately the same energy as the central core (***Durnin et al. (1988))*** and it is the optical energy stored in the concentric rings that is capable of regenerating the beam profile if the central lobe is obstructed ***(MacDonald et al. (1996),*** Figure ***Box*** 5-b), however, the same optical energy is also capable of generating out-of-focus fluorescence. In general, images acquired with a Bessel beam therefore exhibit lower contrast and inferior optical sectioning compared to Gaussian illumination, though confocal line detection ***(Fahrbach and Rohrbach (2012))*** ammeliorates signal to background.

Bessel beams are robust to imaging in scattering media ***(Fahrbach et al. (2010))*** and although this “self-healing” has been well studied in the past, the application of Bessel beams to LSM has been so far more geared towards isotropic resolution ***(Planchon et al. (2011)***; ***Gao et al. (2014))*** or studies of the scattering properties of the sample and the self-reconstructing properties of the Bessel beam itself ***(Fahrbach et al. (2013, 2010))***. Finally, two groups have published technological development concerning Bessel beams applied to LSM ***(Zhang et al. (2014); Zhao et al. (2016)).***

**Figure 6.**
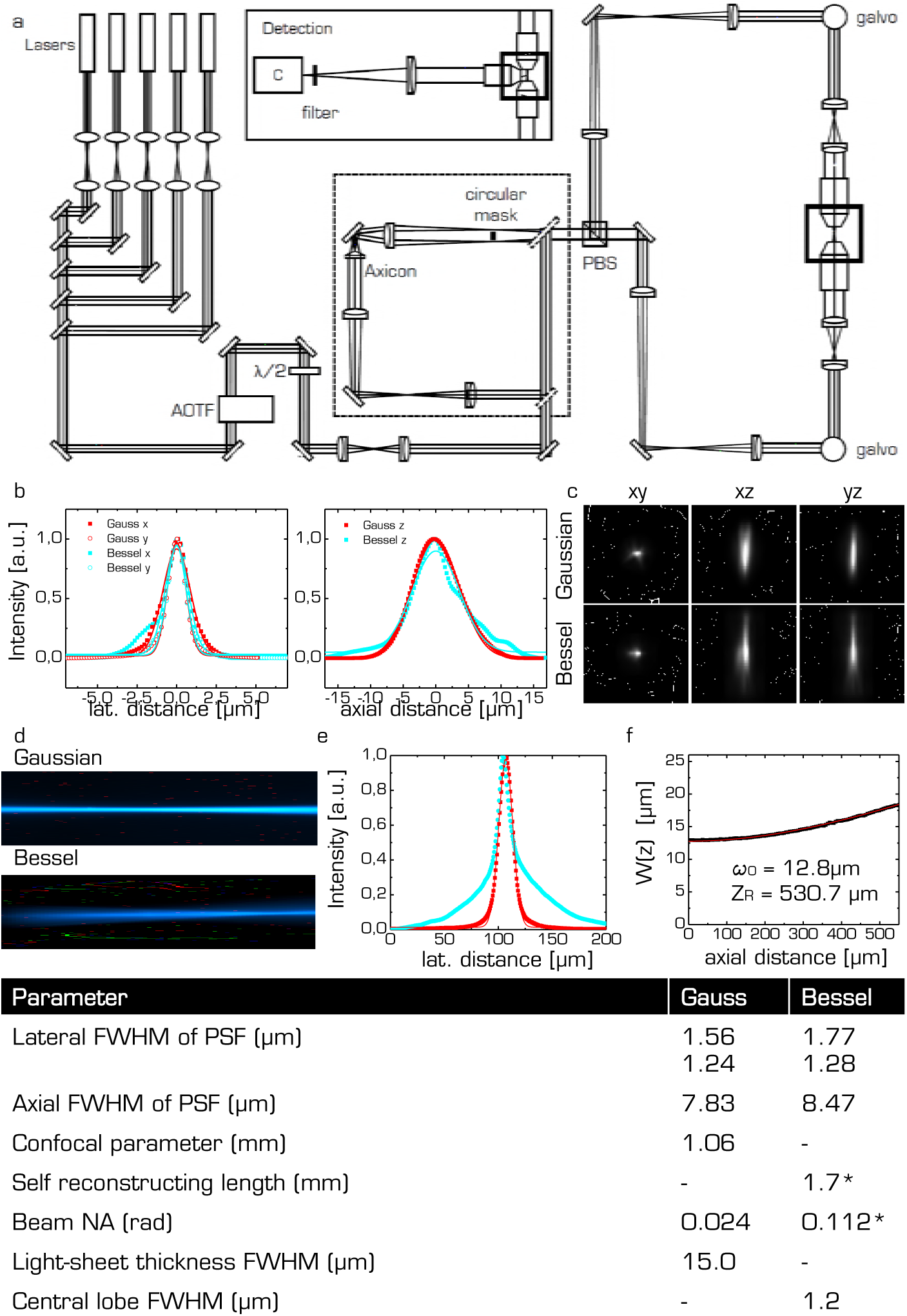
**a)** The custom-made LSM uses double sided illumination by either Gaussian or Bessel beams. Lasers are multiplexed into a single excitation line and an acousto-optical tunable filter (AOTF) is used to select the laser line and regulate power. A half wave plate (λ/2) is used to split the excitation into one of the two equivalent excitation arms using a polarizing beam splitter (PBS). Two flip mirrors allow the user to switch between Gaussian and Bessel beam illumination path (dashed box) which contained the axicon. The light sheets are created digitally by scanning to galvanometric mirrors (galvos) and projected into the sample chamber by two excitation objectives (10x, 0.3NA, WD 17.5 mm). Detection occurs with a specialized objective (10x,0.6NA, WD 12 mm) in a simple wide-field scheme (inset) using a tube lens and a sCMOS camera (C) with confocal line read out mode. See methods for details. **b)** Measured point spread functions (PSFs) for the lateral and axial direction for Gaussian (red) and Bessel illumination (cyan) using fluorescent beads of 100 nm diameter. Beads were automatically extracted and distilled into a PSF using commercial software (PSF distiller, Huygens Software, Scientific Volume Imaging BV, Hilversum, The Netherlands). The full width half maximum (FWHM) of the PSF are reported in the table for Gaussian and Bessel illumination respectively. **c)** PSFs for Gaussian and Bessel illumination. **d)** Longitudinal beam profile. **e)** Transversal profile for the Gaussian (red) and Bessel beam (cyan). f) Beam width ω(z) of the Gaussian beam extracted from profile shown in **(f)**. Red line indicates fit to hyperbolic function. The beam waist ω_0_, the Rayleigh range ***Z_R_*** and the beam NA were extracted from the fit. Bottom: table of all beam parameters. * indicates theoretically derived values.

**Figure 7.**
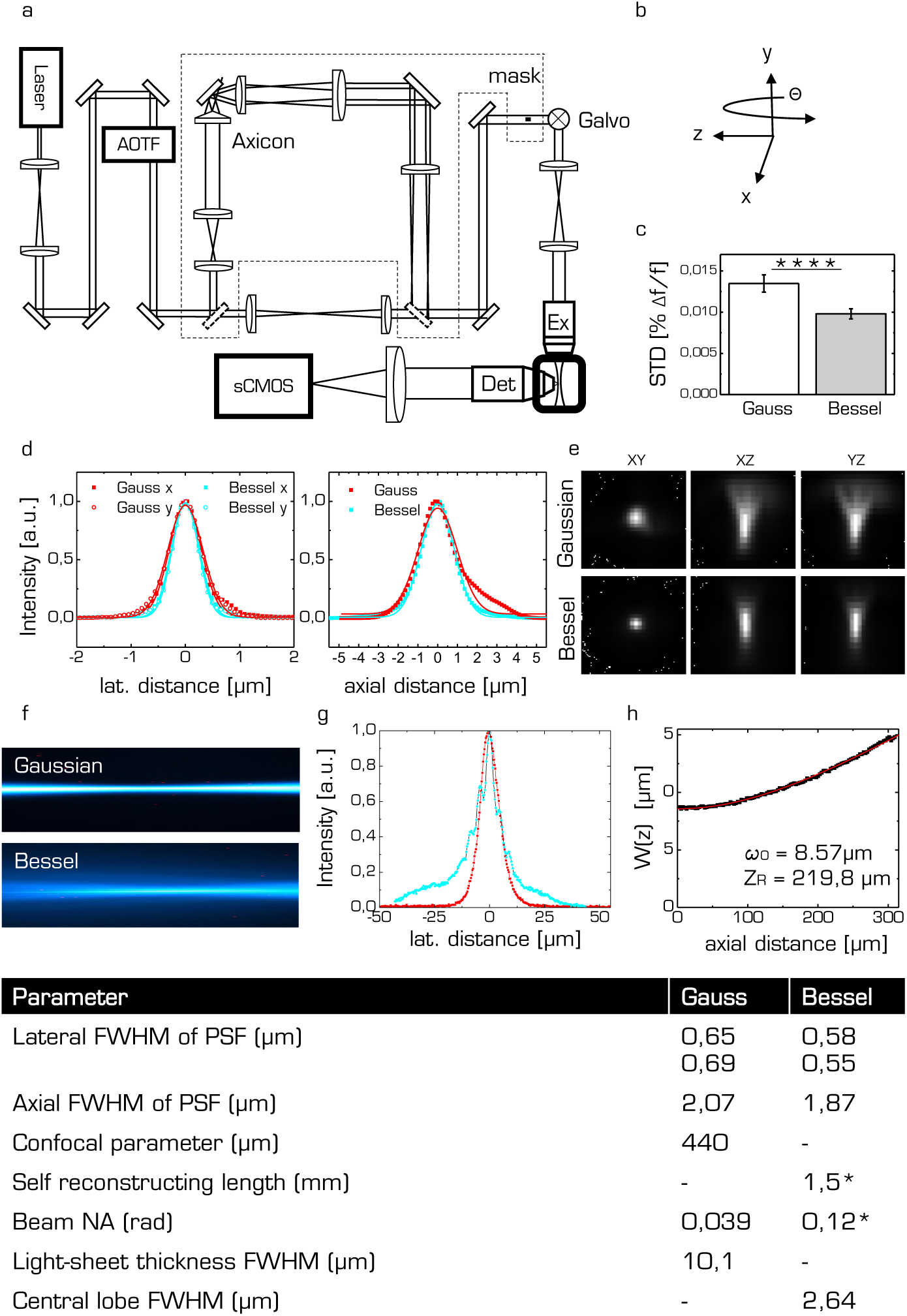
Light-sheet microscope for functional imaging. **a)** A diode laser at 488 nm was used to excite GCaMP6f and an acousto optical tunable filter (AOTF) was used to regulate power. The light sheet was created digitally with a galvanometric mirror (galvo) which was re-imaged onto the back focal plane of the excitation objective (4x, 0.13 NA, air immersion, Olympus, Tokyo, Japan). Fluorescence was detected by a water immersion objective (20x, 1 NA, water immersion, Olympus, Tokyo, Japan) and imaged with a tube lens onto the chip of a fast sCMOS camera. Flip mirrors were used to direct the excitation light into an alternative path with an axicon to generate a Bessel beam. **b)** Sample mounting: zebrafish larvae were embedded in low-melting point agarose and drawn with a syringe into a glass capillary. The capillary was mounted onto an x-, y-, z-, Θ-stage. **c)**. Standard deviation of the flickering measured with fluorescein in the sample chamber and by selecting ROIs the typical size of cells evenly distributed over the field of view. **d)** Measured point spread functions (PSFs) for the lateral and axial directions for Gaussian (red) and Bessel illumination (cyan) using fluorescent beads of 100 nm diameter. Beads were automatically extracted and distilled into a PSF using commercial software (PSF distiller, Huygens Software, Scientific Volume Imaging BV, Hilversum, The Netherlands). The full width half maximum (FWHM) of the PSF are reported in the table for Gaussian and Bessel illumination respectively. **e)** PSFs for Gaussian and Bessel illumination. **f)** Longitudinal beam profile. **g)** Transversal profile for the Gaussian (red) and Bessel beam (cyan). **h)** Beam width ω(z) of the Gaussian beam extracted from profile shown in (f). Red line indicates fit to hyperbolic function. The beam waist ω_**0**_ and the Rayleigh range ***Z_R_*** were extracted from the fit. Bottom: table of all beam parameters. * indicates theoretically derived values.

**Figure 8.**
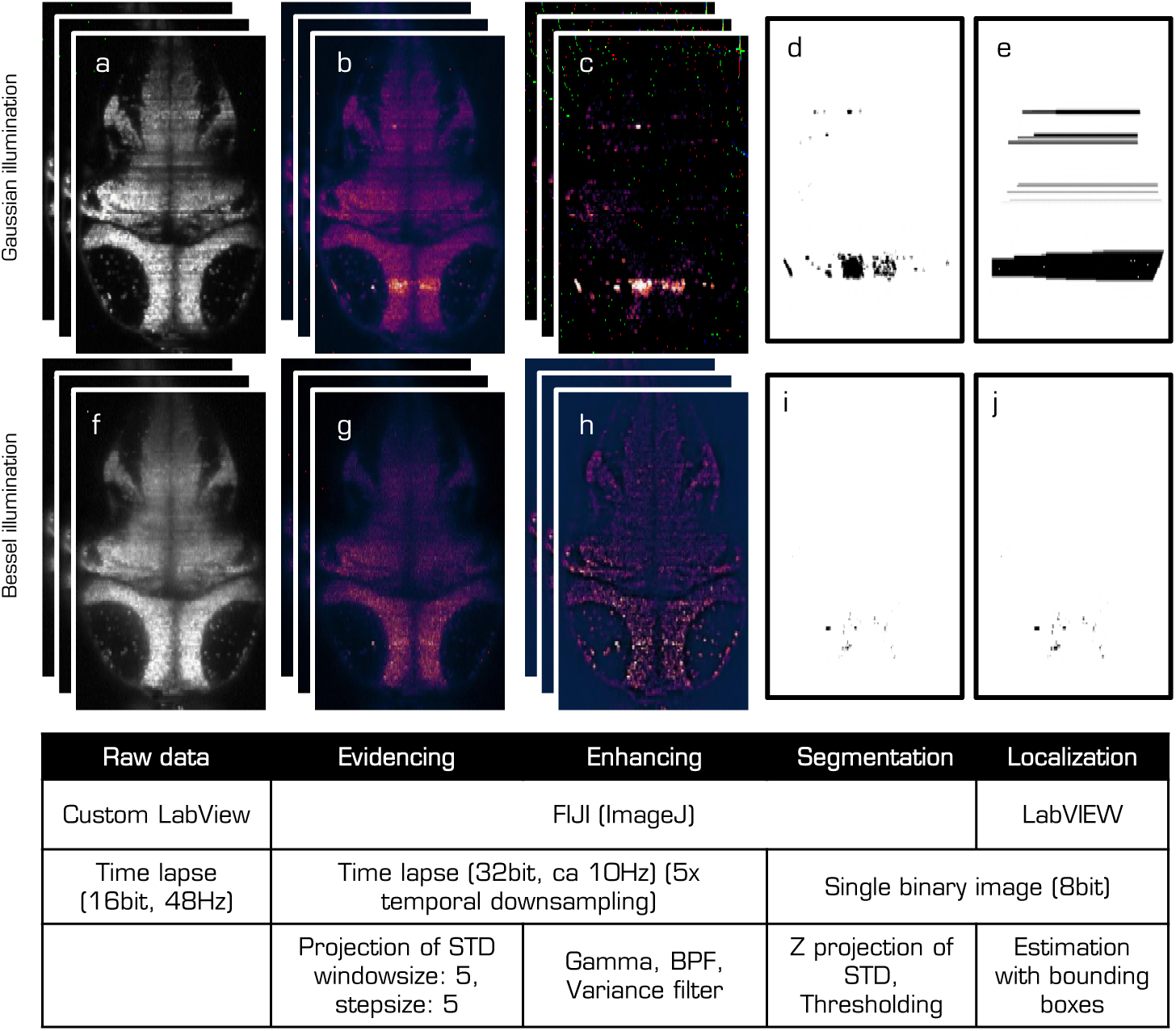
Work flow to estimate area severely affected by primary flickering. The first row refers to data obtained with Gaussian illumination while the second row was obtained with Bessel illumination. The raw data stack **(a,f)** was first temporally downsampled by calculating projections of the STD over 5 frame windows. Those STD stacks **(b,g)** were then treated over the entire image to further enhance the flickering areas, evidenced as bright horizontal stripes, using Gamma, Band path filter (BPF) and variance filter. The resulting time lapses **(c,h)** are projected along z to create single STD images, which were adjusted to the same brightness scale before a common threshold was applied to binarize them **(d,i)**. The binary images were then evaluated with a custom-written LabVIEW program to automatically localize the “blobs”. The bounding boxes (e,j) were used to calculate the overall area affected by flickering. See methods for details. **Figure 3-supplementary Movie 1.** Raw data time lapse imaged with a Gaussian beam (a, supplementary movie **20**) **Figure 3-supplementary Movie 2.** Raw data time lapse imaged with a Bessel beam (f, supplementary movie **21**) **Figure 3-supplementary Movie 3.** Standard deviation time lapse imaged with a Gaussian beam (b, supplementary movie **22**) **Figure 3-supplementary Movie 4.** Standard deviation time lapse imaged with a Bessel beam (g, supplementary movie 23)

**Figure 9.**
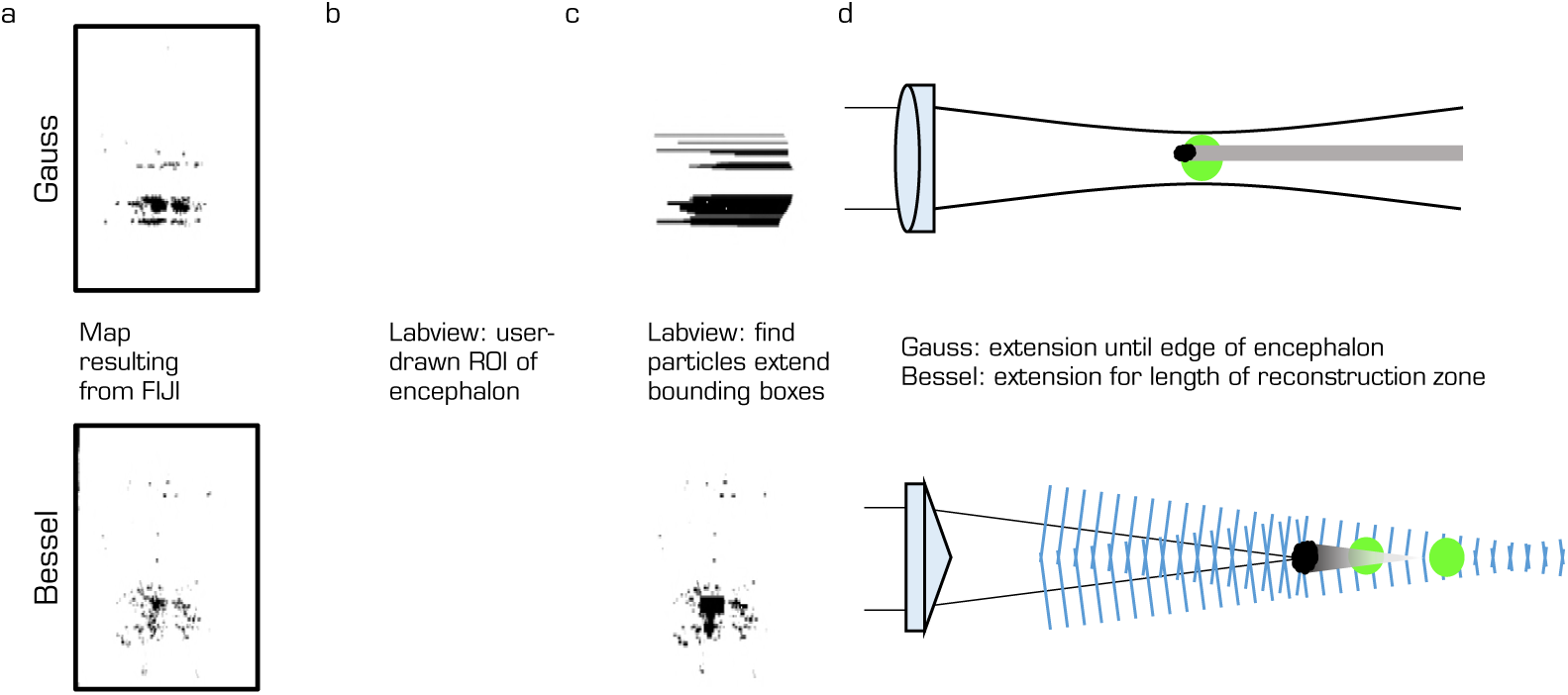
Extension of bounding boxes. The upper row refers to Gaussian illumination whereas the lower row refers to Bessel beam illumination. The binary image **(a)** obtained from the methodology illustrated in ***Figure 8*** was loaded into a custom-written LabVIEW program that automatically determined the bounding boxes of each identified particle. The only input of the user was a free-hand selection of the larva encephalon **(b)** that was used for both Bessel and Gauss evaluation. For Gaussian illumination the bounding boxes were horizontally extruded until they reached the limits of the encephalon mask **(c)** whereas for Bessel illumination the bounding box were extended up to a length corresponding to the reconstruction length of the Bessel beam in the sample plane ***(Equation 1***). The LabVIEW program then automatically summed the area of all bounding boxes and their percentage of the entire encephalon. **(d)** Schematic on the casting of shadows for Gaussian and Bessel beam illumination.

**Figure 10.**
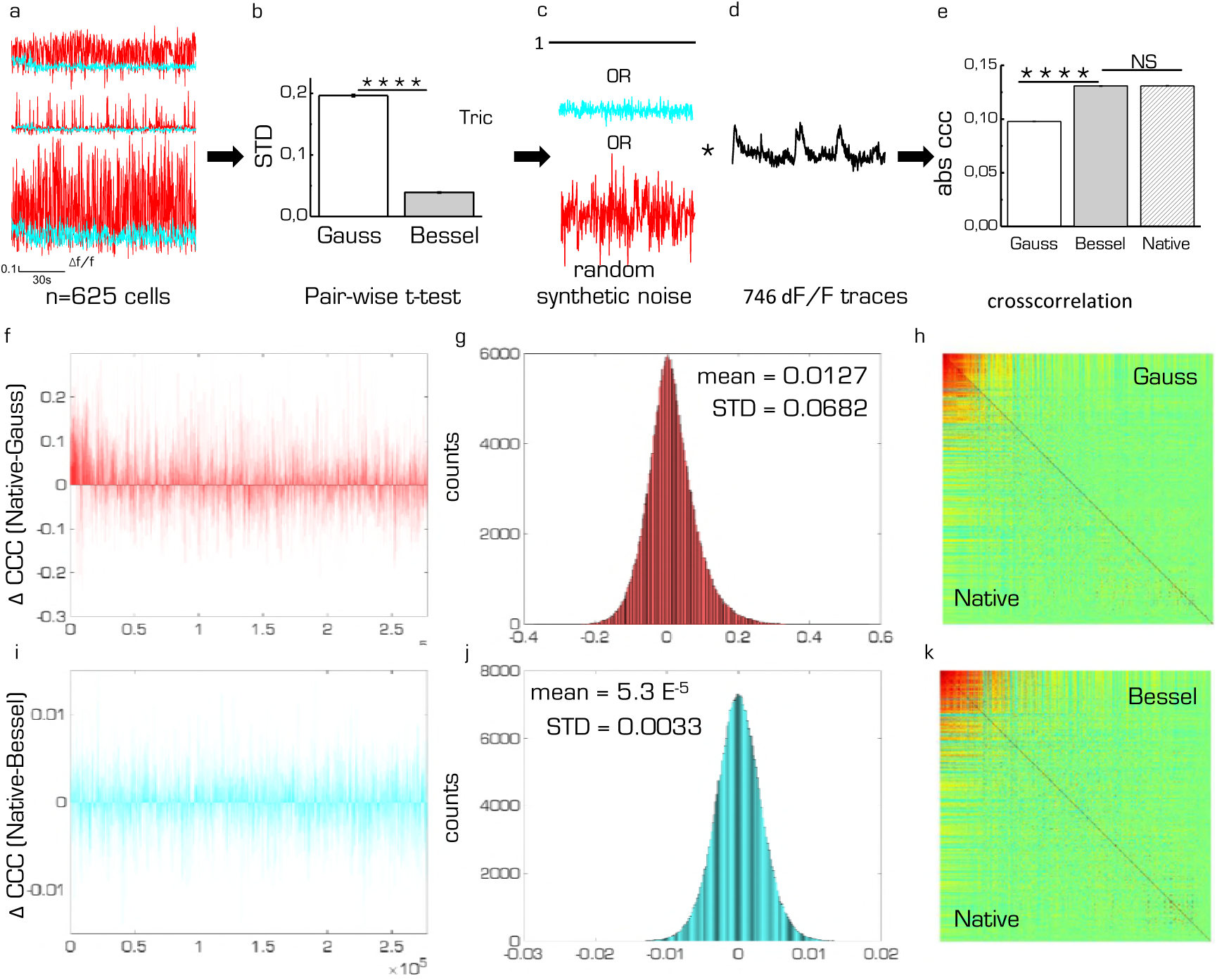
Correlation matrix showing the effect of random synthetic noise. The conceptual flow is as follows: **(a)** from n=625 cells with inhibited activity (Tricaine) an average baseline noise was calculated using a pair-wise t-test for Gaussian and Bessel beam illumination respectively **(b)**. This noise level was used as amplitude to generate random white noise **(c)** which was point-wise multiplied to 746 dF/F traces measured with Bessel beam modality **(d)**. By defining the 746 synthetic noise-free traces as ground truth (native), we obtained three data sets that we compared by respective cross correlation. **(e)** Average absolute cross correlation coefficient (ccc) (*p* < 0.0001, t-test, n= 277885 corresponding to 746 cells, error is sem, NS: not significant). To obtain a measure of accuracy, we subtracted the Gaussian cross correlation coefficients from the native counterparts **(f)**. The histogram of this distribution **(g)** yielded a mean indicating a bias of a measurement made in Gaussian modality, whereas the STD is a measure for its accuracy. **(h)** Correlation matrix showing the native traces (without synthetic noise) in the lower triangular matrix and the traces with multiplied Gaussian noise in the upper triangular matrix. **(i,j,k))** Same as f,g,h) but for Bessel beam modality.

**Figure 11.**
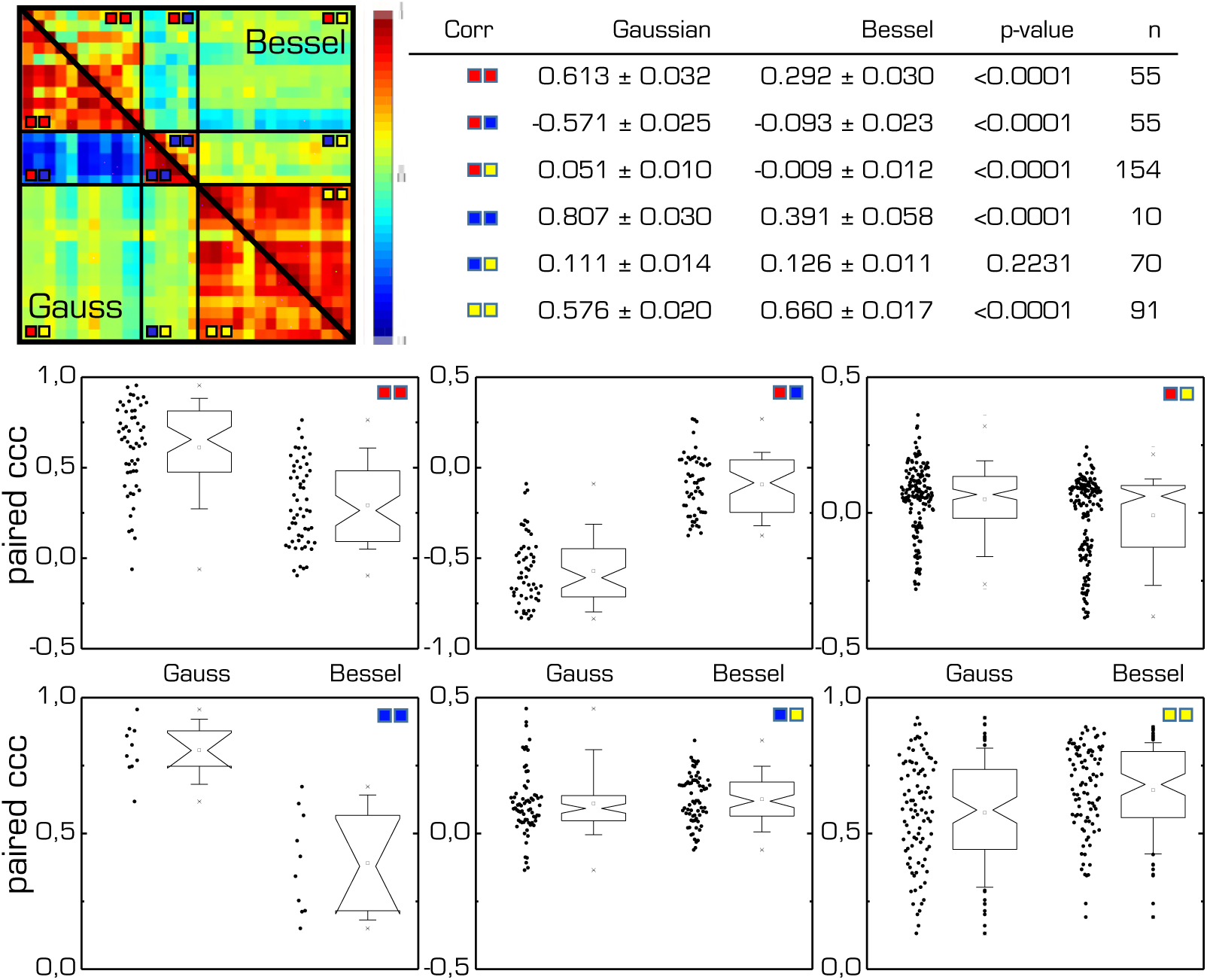
Comparison of the averaged cross correlation coefficients for groups of cells for Gaussian and Bessel beam illumination. Colors correspond to groups of cells marked in ***Figure 4-i)***. Table lists mean of paired t-test, error is sem. Box plots show corresponding data points.

**Figure 12.**
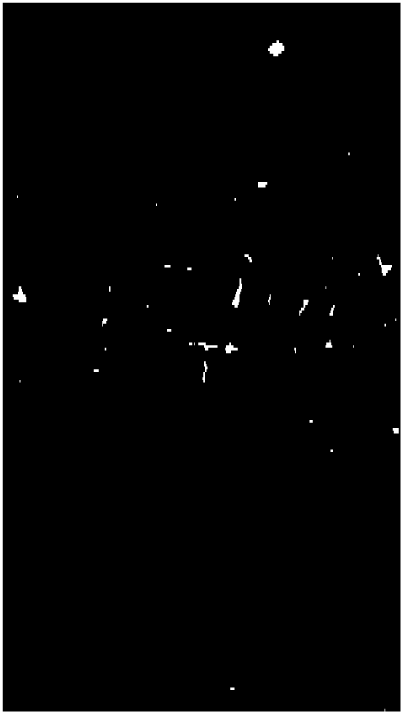
Raw data of the vasculature of a Thy1-GFP-M mouse labelled with tetramethylrhodamine-albumin imaged with Gaussian illumination. See Main ***Figure 2*** for details.

**Figure 13.**
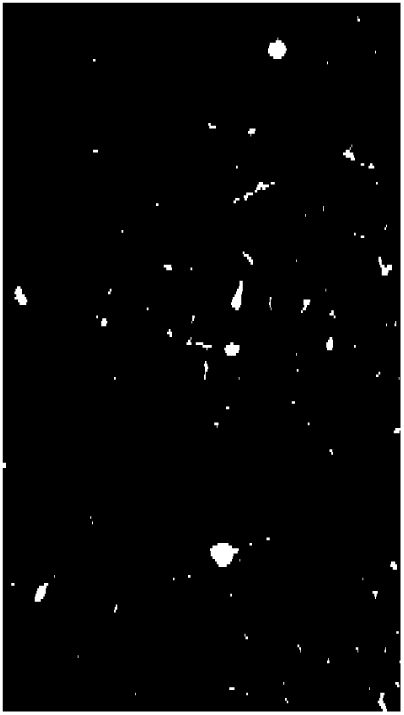
Raw data of the vasculature of a Thy1-GFP-M mouse labelled with tetramethylrhodamine-albumin imaged with Bessel beam illumination. See Main ***Figure 2*** for details.

**Figure 14.**
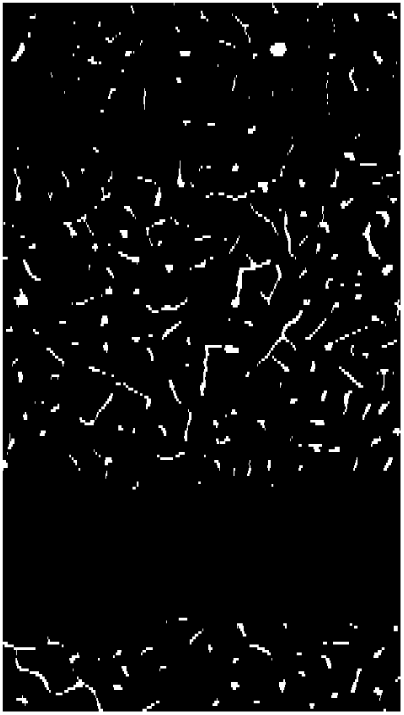
Segmented data of the vasculature of a Thy1-GFP-M mouse labelled with tetramethylrhodamine-albumin imaged with Gaussian illumination. Automated segmentation was based on simple thresholding. See Main ***Figure 2*** for details.

**Figure 15.**
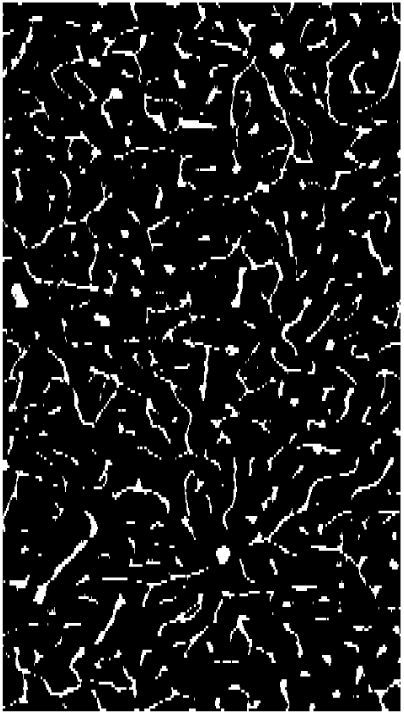
Segmented data of the vasculature of a Thy1-GFP-M mouse labelled with tetramethylrhodamine-albumin imaged with Bessel beam illumination. Automated segmentation was based on simple thresholding. See Main ***Figure 2*** for details.

**Figure 16.**
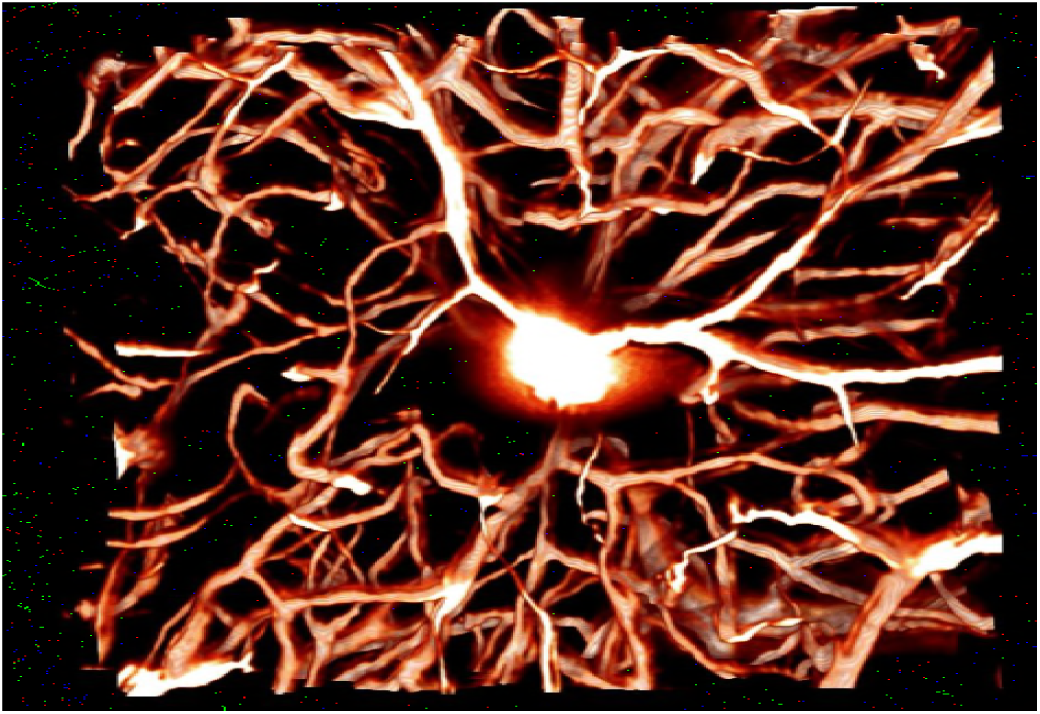
3D projection of raw data of the vasculature of a Thy1-GFP-M mouse labelled with tetramethylrhodamine-albumin imaged with Gaussian beam illumination. A look-up table was applied for clarity. See Main ***Figure2*** for details.

**Figure 17.**
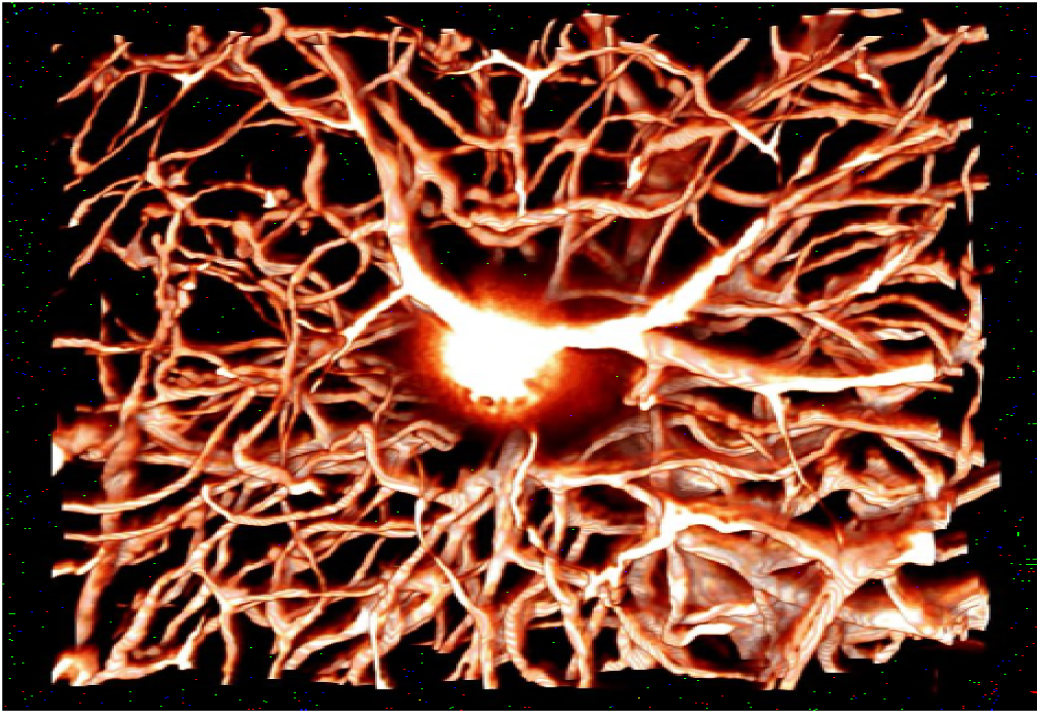
3D projection of raw data of the vasculature of a Thy1-GFP-M mouse labelled with tetramethylrhodamine-albumin imaged with Bessel beam illumination. A look-up table was applied for clarity. See Main ***Figure2*** for details.

**Figure 18.**
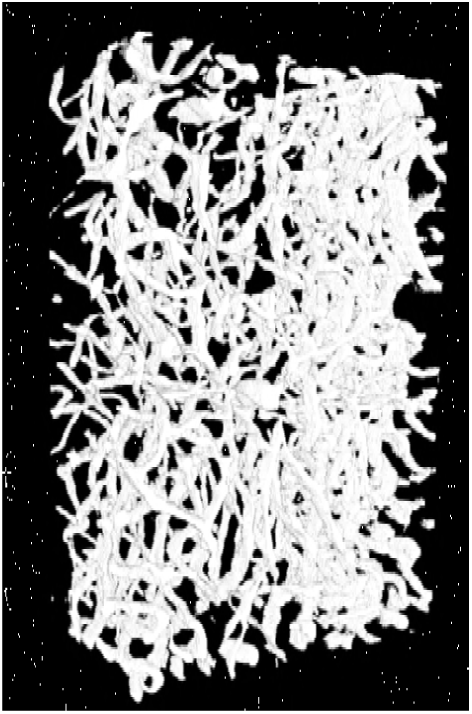
3D projection of segmented data of the vasculature of a Thy1-GFP-M mouse labelled with tetramethylrhodamine-albumin imaged with Gaussian beam illumination. Automated segmentation was based on simple thresholding. See Main *Figure 2* for details.

**Figure 19.**
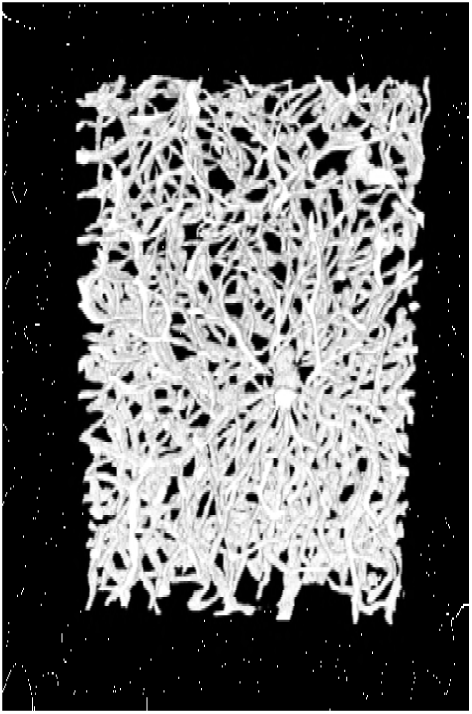
3D projection of segmented data of the vasculature of a Thy1-GFP-M mouse labelled with tetramethylrhodamine-albumin imaged with Bessel beam illumination. Automated segmentation was based on simple thresholding. See Main ***Figure 2*** for details.

**Figure 20.**
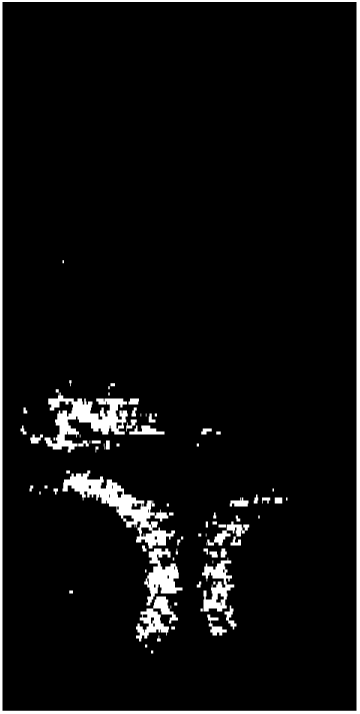
Transverse plane of a 5dpf elavl3:H2B-GCaMP6s larva. Raw data time lapse imaged with a Gaussian beam. See supplementary ***Figure 8*** for details.

**Figure 21.**
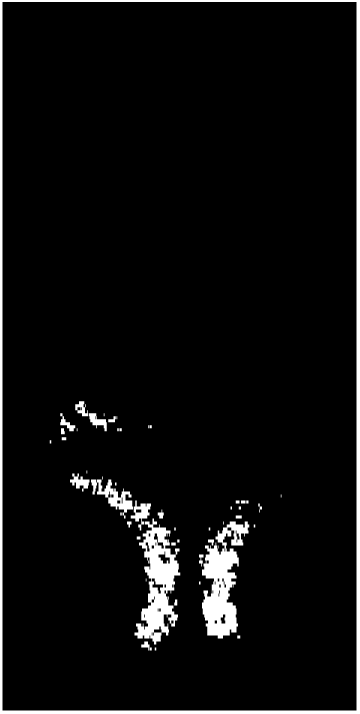
Transverse plane of a 5dpf elavl3:H2B-GCaMP6s larva. Raw data time lapse imaged with a Bessel beam. See supplementary ***Figure 8*** for details.

**Figure 22.**
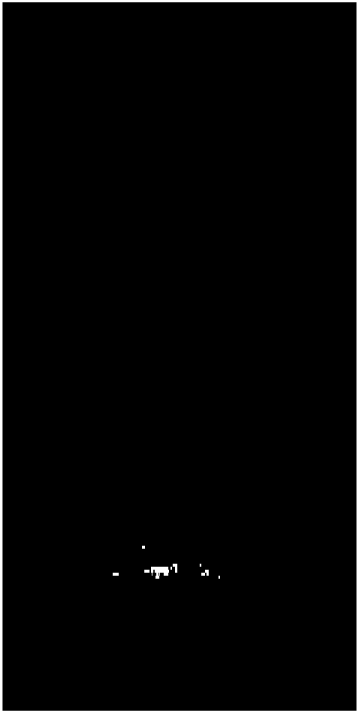
Transverse plane of a 5dpf elavl3:H2B-GCaMP6s larva. Standard deviation time lapse imaged with a Gaussian beam. See supplementary ***Figure 8*** for details.

**Figure 23.**
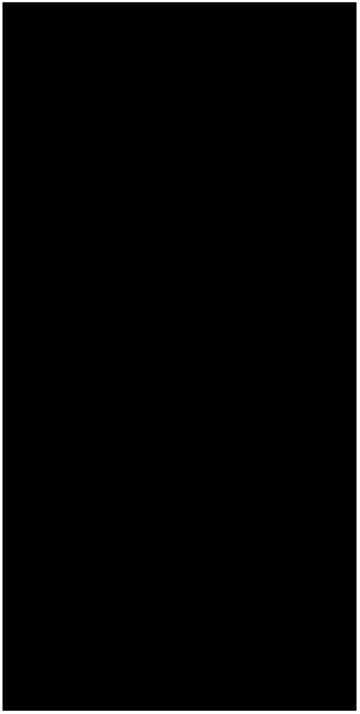
Transverse plane of a 5dpf elavl3:H2B-GCaMP6s larva. Standard deviation time lapse imaged with a Bessel beam. See supplementary ***Figure 8*** for details.

**Figure 24.**
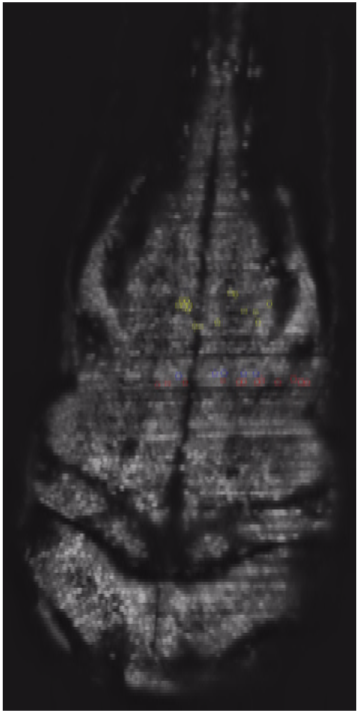
Transverse plane of a 4dpf elavl3:H2B-GCaMP6s larva. Time lapse of spontaneous neuronal activity imaged with a Gaussian beam. See main Figure 4 for details. The raw data was despeckled and local contrast was enhanced with standard FIJI functions.

**Figure 25.**
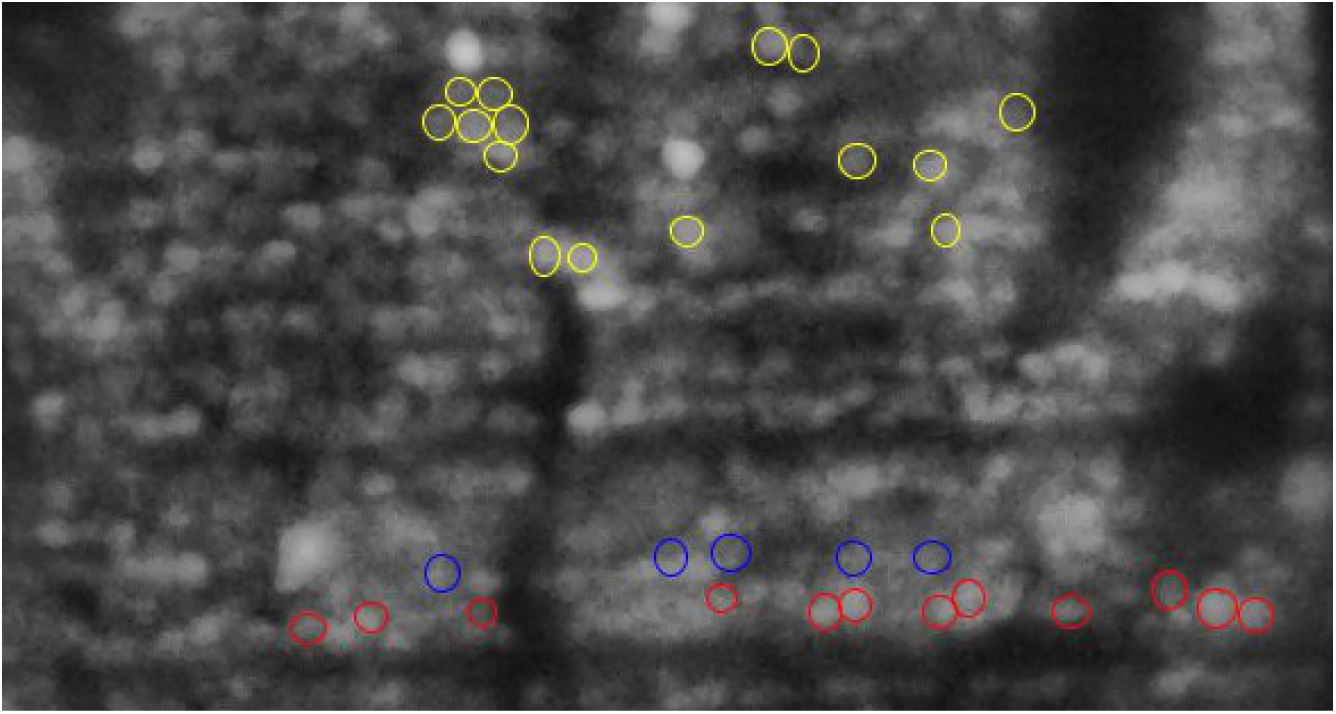
Transverse plane of a 4dpf elavl3:H2B-GCaMP6s larva. Time lapse detailing the cells of spontaneous neuronal activity imaged with a Gaussian beam. See main ***Figure 4*** for details. The raw data was despeckled and local contrast was enhanced with standard FIJI functions.

## References

Acciai L, Soda P, Iannello G. Automated neuron tracing methods: an updated account. Neuroinformatics. 2016; 14(4):353–367.

Ahrens MB, Li JM, Orger MB, Robson DN, Schier AF, Engert F, Portugues R. Brain-wide neuronal dynamics during motor adaptation in zebrafish. Nature. 2012; 485(7399):471–477.

Attili S, Hughes SM. Anaesthetic tricaine acts preferentially on Neural voltage-gated sodium channels and fails to block directly evoked muscle contraction. PloS one. 2014; 9(8):e103751.

Attwell D, Buchan AM, Charpak S, Lauritzen M, MacVicar BA, Newman EA. Glial and neuronal control of brain blood flow. Nature. 2010; 468(7321):232–243.

Bria A, Iannello G. TeraStitcher-a tool for fast automatic 3D-stitching of teravoxel-sized microscopy images. BMC bioinformatics. 2012; 13(1):316.

Bucur O, Irshad H, Montaser-Kouhsari L, Knoblauch NW, Oh EY, Nowak J, Beck AH, 3D morphological hallmarks of breast carcinogenesis: Diagnosis of non-invasive and invasive breast cancer with Lightsheet microscopy. AACR; 2015.

Chen Y, Glaser A, Liu JT. Bessel-beam illumination in dual-axis confocal microscopy mitigates resolution degradation caused by refractive heterogeneities. Journal of Biophotonics. 2016;.

Chitalia R, Mueller J, Fu HL, Whitley MJ, Kirsch DG, Brown JQ, Willett R, Ramanujam N. Algorithms for differentiating between images of heterogeneous tissue across fluorescence microscopes. Biomedical Optics Express. 2016; 7(9):3412–3424.

Chung K, Wallace J, Kim SY, Kalyanasundaram S, Andalman AS, Davidson TJ, Mirzabekov JJ, Zalocusky KA, Mattis J, Denisin AK, et al. Structural and molecular interrogation of intact biological systems. Nature. 2013; 497(7449):332–337.

Costantini I, Ghobril JP, Di Giovanna AP, Mascaro ALA, Silvestri L, Müllenbroich MC, Onofri L, Conti V, Vanzi F, Sacconi L, et al. A versatile clearing agent for multi-modal brain imaging. arXiv preprint arXiv:150403855. 2015;.

Dodt HU, Leischner U, Schierloh A, Jährling N, Mauch CP, Deininger K, Deussing JM, Eder M, Zieglgänsberger W, Becker K. Ultramicroscopy: three-dimensional visualization of neuronal networks in the whole mouse brain. Nature methods. 2007; 4(4):331.

Durnin J, Eberly J, Miceli J. Comparison of Bessel and Gaussian beams. Optics letters. 1988; 13(2):79–80.

Durnin J, Miceli Jr J, Eberly J. Diffraction-free beams. Physical review letters. 1987; 58(15):1499.

Fahrbach FO, Gurchenkov V, Alessandri K, Nassoy P, Rohrbach A. Light-sheet microscopy in thick media using scanned Bessel beams and two-photon fluorescence excitation. Optics express. 2013; 21(11):13824–13839.

Fahrbach FO, Rohrbach A. Propagation stability of self-reconstructing Bessel beams enables contrast-enhanced imaging in thick media. Nature communications. 2012; 3:632.

Fahrbach FO, Simon P, Rohrbach A. Microscopy with self-reconstructing beams. Nature Photonics. 2010; 4(11):780–785.

Frasconi P, Silvestri L, Soda P, Cortini R, Pavone FS, Iannello G. Large-scale automated identification of mouse brain cells in confocal light sheet microscopy images. Bioinformatics. 2014; 30(17):i587–i593.

Freeman J, Vladimirov N, Kawashima T, Mu Y, Sofroniew NJ, Bennett DV, Rosen J, Yang CT, Looger LL, Ahrens MB. Mapping brain activity at scale with cluster computing. Nature methods. 2014; 11(9):941–950.

Gao L, Shao L, Chen BC, Betzig E. 3D live fluorescence imaging of cellular dynamics using Bessel beam plane illumination microscopy. Nature protocols. 2014; 9(5):1083–1101.

Huisken J, Swoger J, Del Bene F, Wittbrodt J, Stelzer EH. Optical sectioning deep inside live embryos by selective plane illumination microscopy. Science. 2004; 305(5686):1007–1009.

Indebetouw G. Nondiffracting optical fields: some remarks on their analysis and synthesis. JOSA A. 1989; 6(1):150–152.

Kawashima T, Zwart MF, Yang CT, Mensh BD, Ahrens MB. The Serotonergic System Tracks the Outcomes of Actions to Mediate Short-Term Motor Learning. Cell. 2016; 167(4):933–946.

Liu S, Zhang D, Liu S, Feng D, Peng H, Cai W. Rivulet: 3d neuron morphology tracing with iterative back-tracking. Neuroinformatics. 2016; 14(4):387–401.

Ma Y, Shaik MA, Kim SH, Kozberg MG, Thibodeaux DN, Zhao HT, Yu H, Hillman EM. Wide-field optical mapping of neural activity and brain haemodynamics: considerations and novel approaches. Phil Trans R Soc B. 2016; 371(1705):20150360.

MacDonald R, Boothroyd S, Okamoto T, Chrostowski J, Syrett B. Interboard optical data distribution by Bessel beam shadowing. Optics Communications. 1996; 122(4-6):169–177.

McGloin D, Dholakia K. Bessel beams: diffraction in a new light. Contemporary Physics. 2005; 46(1):15–28.

Müllenbroich MC, Silvestri L, Onofri L, Costantini I, van’t Hoff M, Sacconi L, Iannello G, Pavone FS. Comprehensive optical and data management infrastructure for high-throughput light-sheet microscopy of whole mouse brains. Neurophotonics. 2015; 2(4):041404–041404.

Pan C, Cai R, Quacquarelli FP, Ghasemigharagoz A, Lourbopoulos A, Matryba P, Plesnila N, Dichgans M, Hellal F, Ertürk A. Shrinkage-mediated imaging of entire organs and organisms using uDISCO. Nature Methods. 2016;.

Panier T, Romano SA, Olive R, Pietri T, Sumbre G, Candelier R, Debrégeas G. Fast functional imaging of multiple brain regions in intact zebrafish larvae using selective plane illumination microscopy. x. 2013; x(x):x.

Peng H, Zhou Z, Meijering E, Zhao T, Ascoli GA, Hawrylycz M. Automatic tracing of ultra-volumes of neuronal images. Nature Methods. 2017; 14(4):332–333.

Planchon TA, Gao L, Milkie DE, Davidson MW, Galbraith JA, Galbraith CG, Betzig E. Rapid three-dimensional isotropic imaging of living cells using Bessel beam plane illumination. Nature methods. 2011; 8(5):417–423.

Richardson DS, Lichtman JW. Clarifying tissue clearing. Cell. 2015; 162(2):246–257.

Rose A, Vision: Human andElectronic. Plenum Press, New YorklLondon; 1973.

Siedentopf H, Zsigmondy R. Uber sichtbarmachung und größenbestimmung ultramikoskopischer teilchen, mit besonderer anwendung auf goldrubingläser. Annalen der Physik. 1902; 315(1):1–39.

Silvestri L, Costantini I, Sacconi L, Pavone FS. Clearing of fixed tissue: a review from a microscopist’s perspective. Journal of biomedical optics. 2016; 21(8):081205–081205.

Susaki EA, Tainaka K, Perrin D, Kishino F, Tawara T, Watanabe TM, Yokoyama C, Onoe H, Eguchi M, Yamaguchi S, et al. Whole-brain imaging with single-cell resolution using chemical cocktails and computational analysis. Cell. 2014; 157(3):726–739.

Tainaka K, Kuno A, Kubota SI, Murakami T, Ueda HR. Chemical principles in tissue clearing and staining protocols for whole-body cell profiling. Annual review of cell and developmental biology. 2016; 32:713–741.

Torres R, Vesuna S, Levene MJ. High-resolution, 2-and 3-dimensional imaging of uncut, unembedded tissue biopsy samples. Archives of Pathology and Laboratory Medicine. 2013; 138(3):395–402.

Tsai PS, Kaufhold JP, Blinder P, Friedman B, Drew PJ, Karten HJ, Lyden PD, Kleinfeld D. Correlations of neuronal and microvascular densities in murine cortex revealed by direct counting and colocalization of nuclei and vessels. Journal of Neuroscience. 2009; 29(46):14553–14570.

Turrini L, Fornetto C, Marchetto G, Müllenbroich M, Tiso N, Vettori A, Resta F, Masi A, Mannaioni G, Pavone F, et al. Optical mapping of neuronal activity during seizures in zebrafish. Scientific Reports. 2017; 7.

Vladimirov N, Mu Y, Kawashima T, Bennett DV, Yang CT, Looger LL, Keller PJ, Freeman J, Ahrens MB. Light-sheet functional imaging in fictively behaving zebrafish. Nature methods. 2014;.

Westerfield M. The zebrafish book: a guide for the laboratory use of zebrafish (Brachydanio rerio). University of Oregon press; 1995.

Zhang P, Phipps M, Goodwin P, Werner J. Confocal line scanning of a Bessel beam for fast 3D Imaging. Optics letters. 2014; 39(12):3682–3685.

Zhao T, Lau SC, Wang Y, Su Y, Wang H, Cheng A, Herrup K, Ip NY, Du S, Loy M. Multicolor 4D Fluorescence Microscopy using Ultrathin Bessel Light Sheets. Scientific reports. 2016; 6.

